# Plastid translocon recycling in dinoflagellates demonstrates the portability of complex plastids between hosts

**DOI:** 10.1101/2024.07.06.602338

**Authors:** William H Lewis, Giulia Paris, Girish Beedessee, Ludek Koreny, Victor Flores, Tom Dendooven, Benoit Gallet, Daniel Yee, Simon Lam, Johan Decelle, Ben F Luisi, Ross F Waller

## Abstract

The plastids of photosynthetic organisms on land are predominantly ‘primary plastids’ derived from an ancient endosymbiosis of a cyanobacterium. Conversely, marine photosynthetic diversity is dominated by plastids gained by subsequent endosymbioses of photosynthetic eukaryotes, so-called ‘complex plastids’. The plastids of major eukaryotic lineages including cryptophytes, haptophytes, stramenopiles, dinoflagellates and apicomplexans, were originally all posited to derive from a single secondary endosymbiosis of a red alga—the ‘chromalveloate’ hypothesis ^1^. Subsequent phylogenetic resolution of eukaryotes indicated that separate events of plastid acquisition must have occurred to account for this distribution of plastids ^2,3^. The number of such events, however, and the donor organisms for the new plastid endosymbioses are still not resolved. A perceived bottleneck of endosymbiotic plastid gain is the development of protein targeting from the hosts into new plastids, and this supposition has often driven hypotheses towards minimising the number of plastid-gain events to explain plastid distribution in eukaryotes. But how plastid protein-targeting is established for new endosymbionts is often unclear, which makes it difficult to assess the likelihood of plastid transfers between lineages. Here we show that Kareniaceae dinoflagellates, that possess complex plastids known to be derived from haptophytes, acquired all the necessary protein import machinery from these haptophytes. Furthermore, cryo-electron tomography revealed that no additional membranes were added to the Kareniaceae complex plastid during serial endosymbiosis, suggesting that the haptophyte-derived import processes were sufficient. Our analyses suggests that complex red plastids are preadapted for horizontal transmission, potentially explaining their widespread distribution in aquatic algal diversity.

## RESULTS AND DISCUSSION

### Haptophyte translocons for all plastid membranes are maintained in Kareniaceae with stable haptophyte-derived plastids

Different Kareniaceae dinoflagellate taxa are known to have different types of plastids. Some have only a peridinin plastid which is the ancestral plastid of dinoflagellates (e.g. *Gertia stigmatica*) ^4^, some continually acquire temporary kleptoplasts from haptophyte prey (e.g. an undescribed taxon, the Ross Sea Dinoflagellate (RSD)) ^5,6^, and others have stable and fully integrated haptophyte-derived plastids (e.g. species of *Karlodinium, Karenia, Takayama*) ^7–9^. These different plastid states seemingly represent a continuum of replacing a photosynthetic peridinin plastid with a new haptophyte-derived plastid to fulfil the role of photosynthesis. The new stable plastids in *Karlodinium, Karenia* and *Takayama* lack any vestige of haptophyte cytosolic organelles or nucleus ^10^, so must rely on protein import from the new host dinoflagellate for their biogenesis and function. Plastids in haptophytes are surrounded by four membranes which require a system of four different translocons for the import of nucleus-encoded proteins for plastid function (**Figure 1A**) ^11^. We asked, how much of this protein import machinery was inherited and redeployed in Kareniaceae dinoflagellates that might have facilitated the integration of these new plastid endosymbionts.

**Figure 1:**
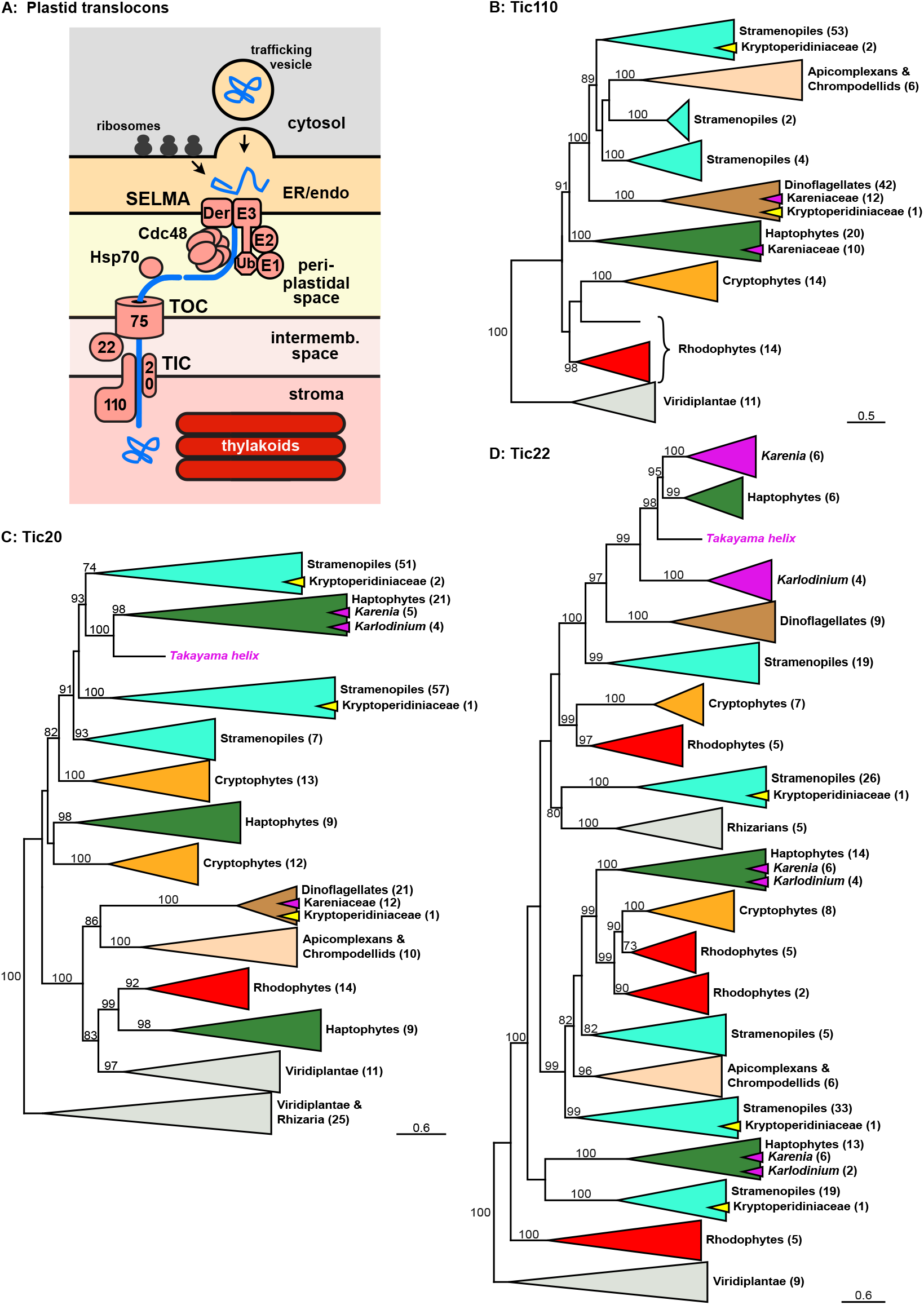
Inner membrane translocons are derived from haptophytes in Kareniaceae. (**A**) Schematic of four-membrane bound plastids and the protein translocation machinery conserved in haptophytes, stramenopiles, cryptophytes and apicomplexans. Protein delivery to the ER or endomembrane system (endo) can occur either via vesicular transport or, in some taxa, co-translational import via ribosomes docked on the outer membrane. The <u>s</u>ymbiont specific <u>E</u>RAD-like <u>ma</u>chinery (SELMA) transfers proteins into the periplastidal space as a relic of the red algal cytoplasm, and TOC and TIC derived from the primary plastid transfer proteins into the plastid stroma. (**BD**) Protein maximum likelihood phylogenies of Tic110, Tic20, and Tic22, respectively. Sequences found in the Kareniaceae monophyletic clades are indicated with magenta triangles or text. Dinotom sequences of the Kryptoperidiniaceae are shown in yellow triangles. Bootstrap support values >70 are shown and the number of sequences per clade are given in parentheses. Scale bars indicate estimate of amino acid substitutions per site. Full phylogenies are given in Supplemental Figure S1.

The inner-most pair of membranes in all known plastids are derived from the primary plastid endosymbiont. These membranes use conserved translocons of the inner and outer chloroplast membranes (TICs and TOCs, respectively) for protein import (**Figure 1A**) ^11^. To test for the presence TIC and TOC components in Kareniaceae, we generated long- and short-read RNA-seq data for *Karlodinium veneficum* and searched this and available transcriptomic data for two further *Karlodinium* spp. (*K. micrum* and *K. armiger*), three *Karenia* spp. (*K. brevis, K. mikimotoi, K. papilionacea*), and *Takayama helix*. Phylogenies were constructed to test for the origin of these proteins. Tic110 and Tic20 are widely conserved inner membrane translocon components and homologues were found throughout the Kareniaceae which grouped specifically with haptophyte orthologues (**Figue 1B-C**, indicated as magenta within green triangles). A major pore-forming translocon element of the TOC complex is Toc75, but as a beta-barrel protein it is often not well conserved and easily recognisable by primary sequence ^12^. We were not able to find homologues of Toc75 in the Kareniaceae, nor any dinoflagellates. However, Tic22 is an intermembrane space protein that acts as a chaperone exchanging proteins from the TOC to TIC ^13^. Multiple paralogues of Tic22 are found throughout plastid types, and all Kareniaceae paralogues were found exclusively within or sister to the haptophyte paralogue clades (**Figue 1D**). Collectively, these data indicate that elements of the haptophyte TIC and TOC complexes have been retained in the Kareniaceae.

The third plastid membrane (counting from the inside) of haptophytes, stramenopiles, cryptophytes and apicomplexans, uses a protein translocon system derived from the endoplasmic reticulum-associated degradation (ERAD) machinery that typically exports misfolded proteins from the ER to the cytoplasm for proteasomal degradation ^14–16^. In an ancestral secondary endosymbiosis of a red alga, most likely in a cryptophyte, the red alga’s ERAD was redeployed from the ER to its relict plasma membrane and repurposed to bring plastid proteins from the host inward into its cytoplasm (**Figure 1A**) ^17,18^. Rather than be degraded, these proteins are then available to the plastid’s TOC and TIC for import into the plastid lumen. This <u>s</u>ymbiont specific <u>E</u>RAD-<u>l</u>ike <u>ma</u>chinery (SELMA) consists of most of the ERAD elements: the derlin component of the membrane translocon, Der1; the ubiquitin-dependent AAA-ATPase Cdc48 that actively translocates the protein cargo; E1, E2 and E3 of the ubiquitination pathway; and chaperones to receive the disordered protein cargo ^14^. The development of this SELMA system to overcome the third membrane was a seminal event in the development of the red algal-derived secondary plastid, enabling essential genes to relocate from the red algal nucleus to that of its new host. Given that ERAD is an essential element of all eukaryotes’ ER, eukaryotes with SELMA contain these proteins as a second paralogous machinery in their third plastid membrane. The molecular phylogenies of Cdc48, Der1 and Uba1 (the E1 ubiquitin activating enzyme) resolve the ERAD proteins separately from SELMA paralogues. The SELMA proteins all group within or sister to the red algal ERAD proteins, consistent with their origin from a red algal-derived secondary plastid (**Figure 2**). Kareniaceae dinoflagellates contain ERAD proteins that group within the rest of dinoflagellates, consistent with the expected vertical inheritance of this essential ER process. Additionally, SELMA paralogues from Kareniaceae dinoflagellates are found, and these group specifically within the haptophyte SELMA clades. In haptophytes, Cdc48 was duplicated and the Kareniaceae dinoflagellates have inherited both Cdc48 SELMA paralogues also (**Figure 2A**). Other ERAD/SELMA components include Ubc4 (the E2 ubiquitin conjugating enzyme), Ubi (the ubiquitin protein itself), and the chaperone Hsp70. These proteins all occur as multiple paralogues in most eukaryotes and the phylogenetic histories of these paralogues are complex. However, all SELMA versions of these proteins in haptophytes have orthologues in *Karlodinium* and *Karenia* that group specifically within these haptophyte clades (**Figure 3**). Therefore, in addition to the TOC/TIC, the haptophyte SELMA protein import machinery for a third plastid membrane has also been retained in the Kareniaceae.

**Figure 2:**
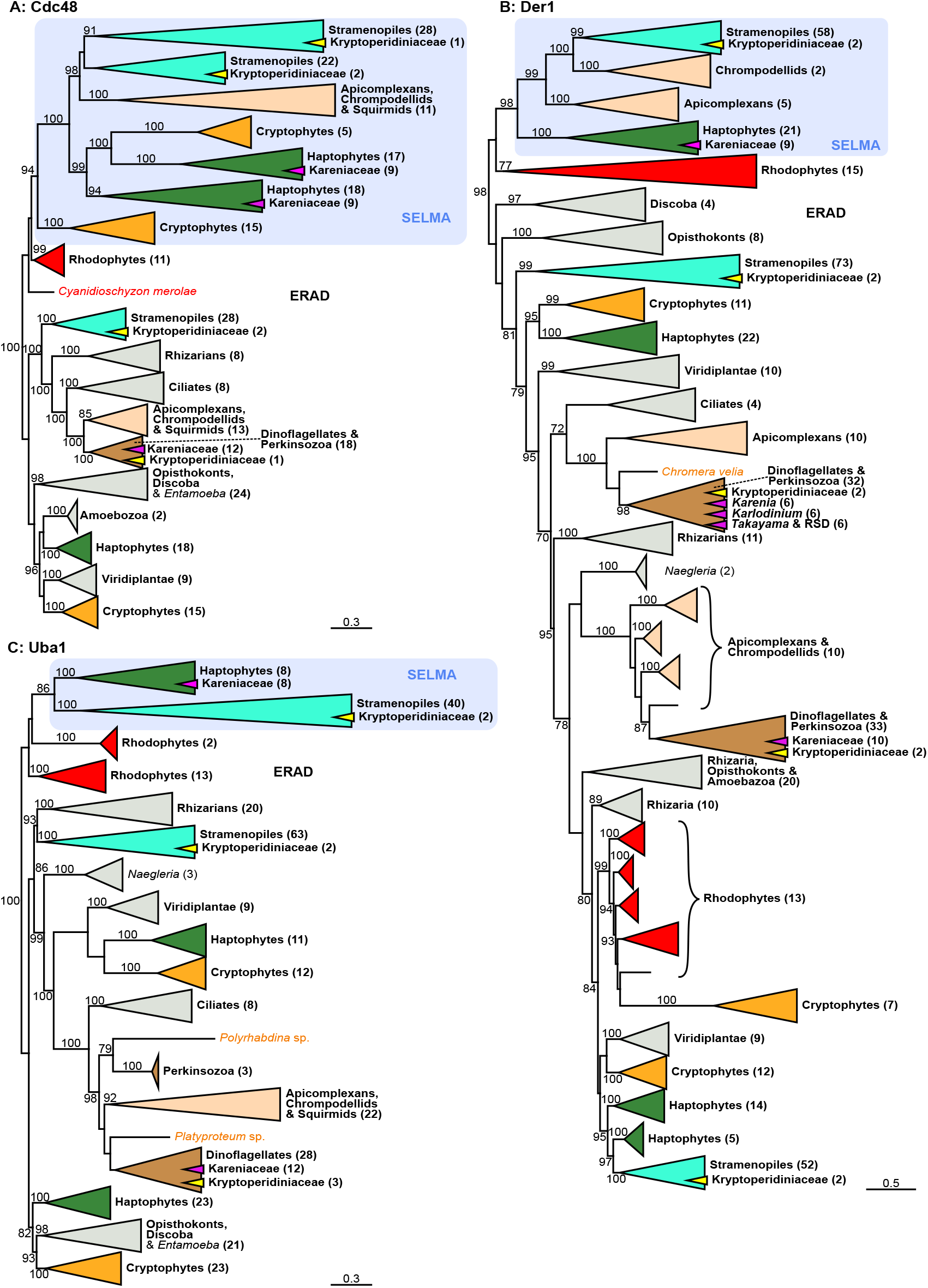
SELMA components Cdc48, Der1 and Uba1 are derived from haptophytes in Kareniaceae. Protein maximum likelihood phylogenies of (**A**) Cdc48, (**B**) Der1, and (**C**) Uba1 shown as for Fig 1. RSD, Ross Sea dinoflagellate which is a kleptoplastic member of the Kareniaceae. Full phylogenies are given in Supplemental Figure S1.

**Figure 3:**
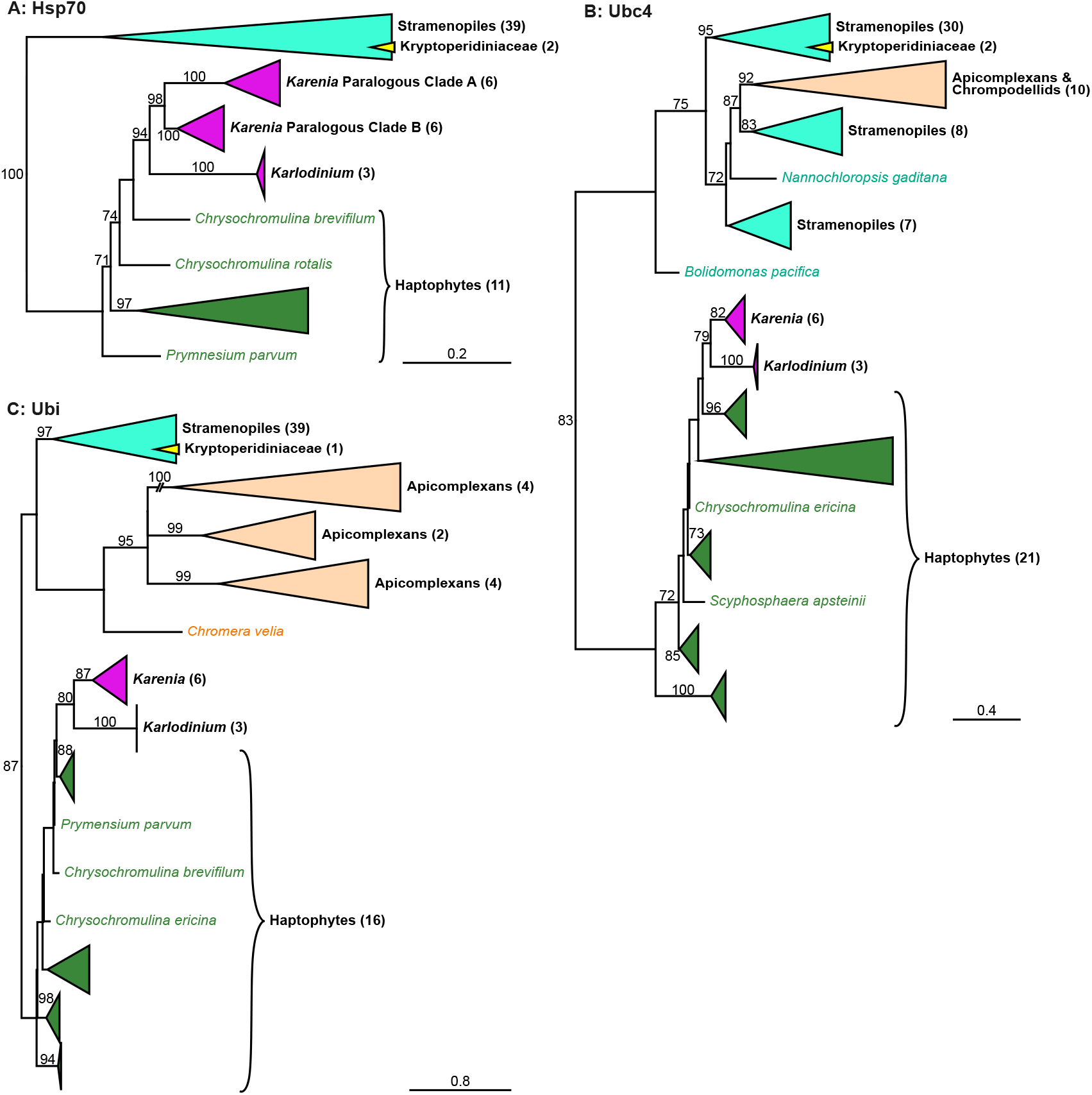
SELMA components Ubc4, Ubi and Hsp70 are derived from haptophytes in Kareniaceae. Protein maximum likelihood phylogenies of SELMA paralogues of (**A**) the chaperone Hsp70, (**B**) the ubiquitin conjugating enzyme Ubc4, and (**C**) ubiquitin (Ubi). are shown as for Fig 1. Full phylogenies are given in Supplemental Figure S1.

The plastids known to use SELMA at the third membrane— in haptophytes, cryptophytes, stramenopiles, and apicomplexans—are surrounded by a fourth, final bounding membrane. This outer membrane is part of the cell’s endomembrane network and either shares direct continuity with the ER or is connected by vesicle trafficking ^1^. Therefore, the first step of protein delivery to these plastids is the cotranslational import of proteins into the ER. Proteins for such plastids bear an N-terminal signal peptide for recognition and delivery into the ER through the Sec61 complex ^19^. This signal peptide is then typically proteolytically removed, and a downstream protein-sorting ‘transit peptide’ is responsible for subsequent recognition and sorting through SELMA, TOC then TIC. To test if the haptophyte-derived SELMA proteins in the Kareniaceae are likely still deployed to and used in plastid import, rather than having been co-opted by its cytosol-orientated ERAD, we examined these proteins for evidence of the bipartite plastid targeting signals. All these SELMA proteins had recognisable signal peptides followed by a further protein extension with the typical features of transit peptides— elevated serine/threonine content, and enrichment for basic over acid residues particularly at the N-terminus (**Figure 4**). These features were also seen for the TIC proteins, and together they support that the haptophyte-derived translocons are all delivered to and function in the plastid.

**Figure 4:**
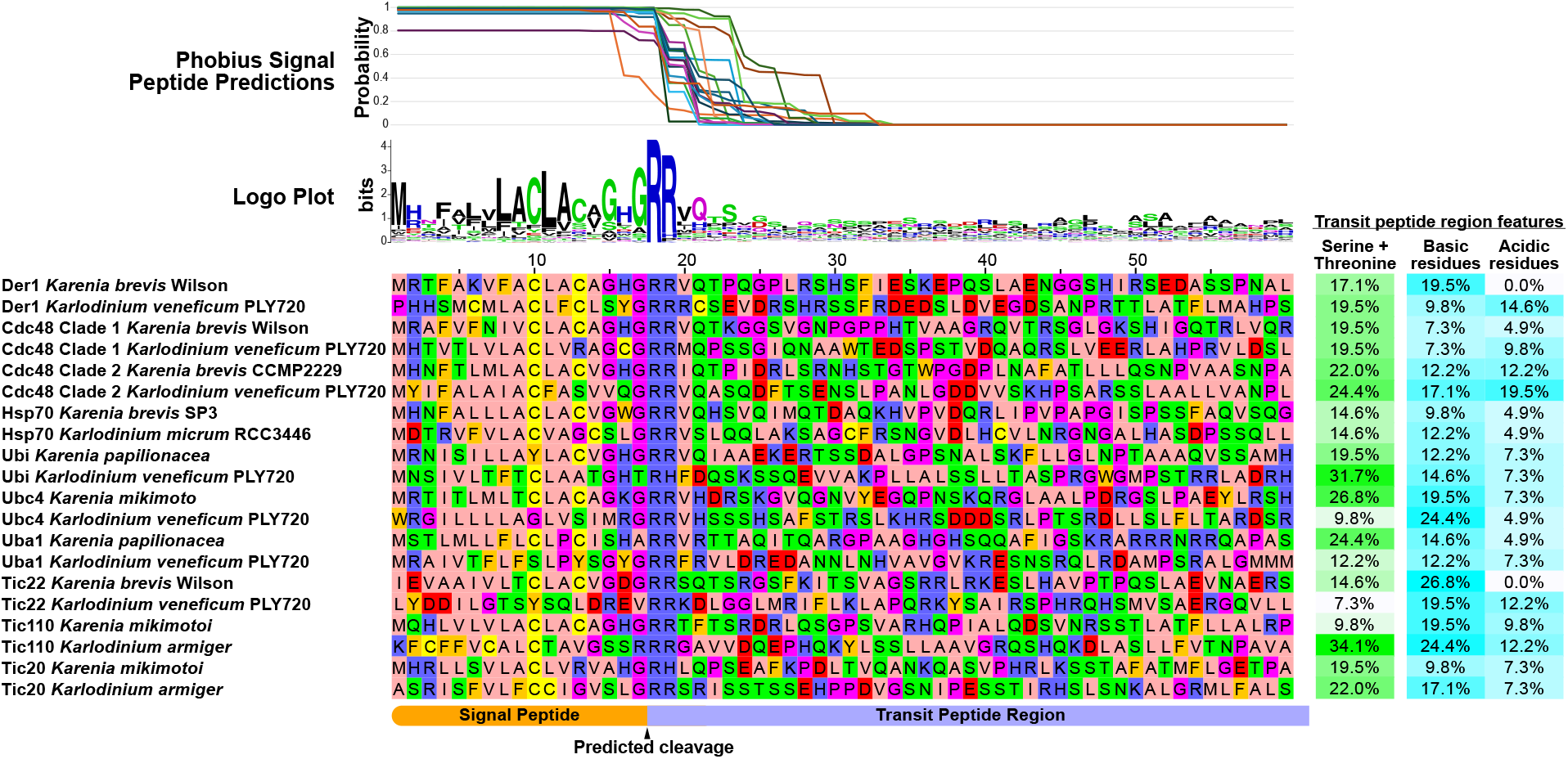
Haptophyte-derived Kareniaceae translocon machinery possess bipartite plastid-targeting presequences. ER-directing signal peptide (SP) predictions by Phobius ^20^ for 20 Kareniaceae translocation components. A logo plot made using WebLogo ^21^ of protein presequences aligned on the predicted cleavage site. Alignment of protein presequences with transit peptide-type features (40 residues post SP cleavage site) shown for each.

### Serial gain of haptophyte plastids in the Kareniaceae did not add further membranes

Our analysis of translocons in the Kareniaceae dinoflagellates accounts for a possible four membranes surrounding these plastids. However, endosymbiotic processes are often associated with the gain of further membranes, typically presumed to derive from the enveloping phagolysosomal membrane and the endosymbiont’s own plasma membrane, although other scenarios of membrane gain have been proposed ^22^. Such acquired, further membranes would presumably require additional protein translocons, which might present further obstacles to endosymbiont establishment. The number of membranes surrounding the plastids of *Karlodinium* and *Karenia* has been previously unclear, with transmission electron microscopy of resin-embedded specimens (**Figure 5A**) consistently showing poor plastid membrane preservation and collapsed membrane profiles. To overcome this, we used cryo-electron tomography (cryoET) on flash frozen *Karlodinium veneficum* cells prepared as thin lamellae cell sections by focused-ion beam (FIB) milling. Reconstructed tomograms of these sections clearly distinguished the separate bounding plastid membranes from the thylakoid membranes and showed the number of bounding membranes to be four (**Figure 5B-C**). These data show that the acquisition of the haptophyte plastid in Kareniaceae dinoflagellates did not result in additional bounding membranes and, therefore, the haptophyte-derived translocons alone are likely sufficient for protein import.

**Figure 5:**
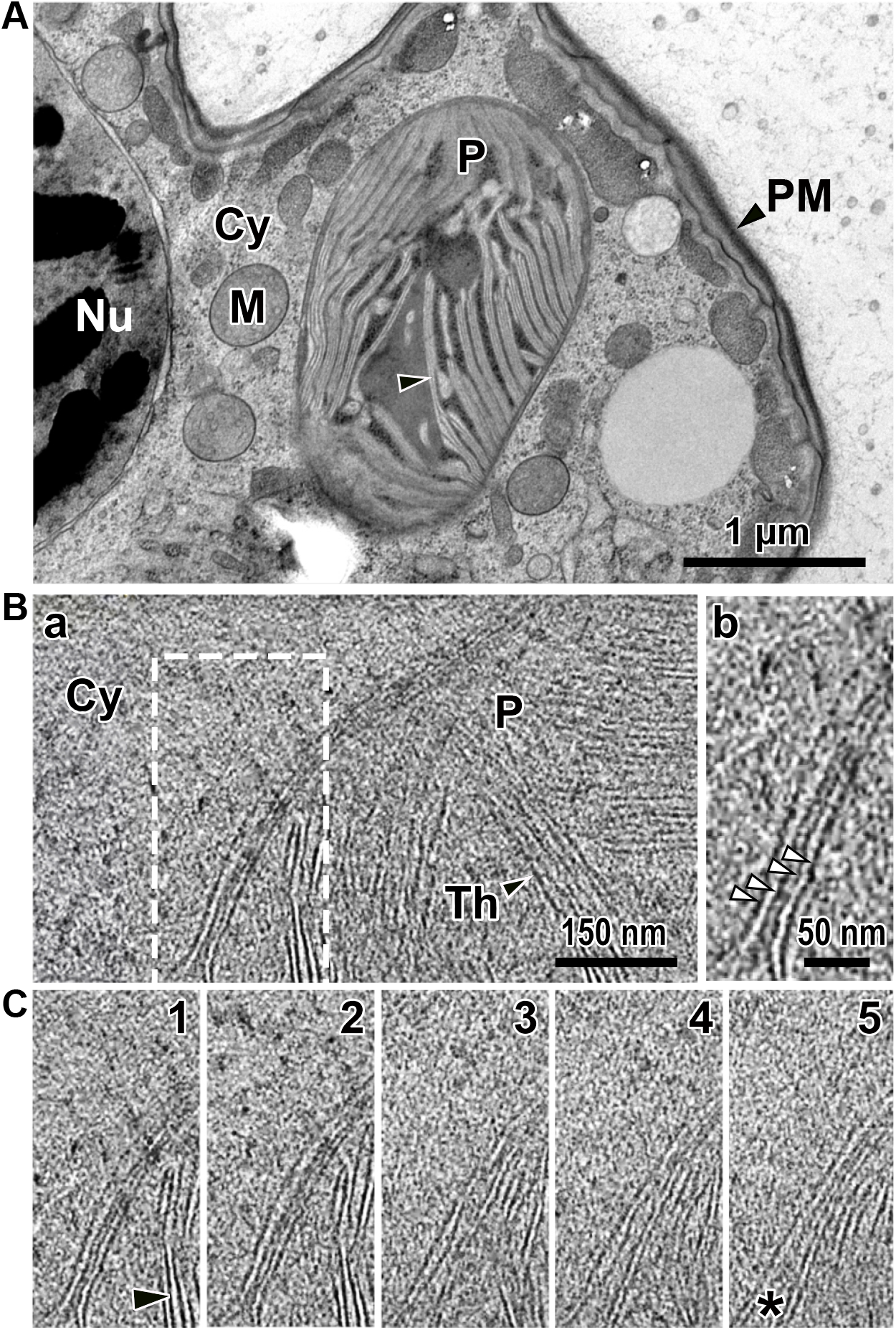
*Karlodinium veneficum* plastids are surrounded by four membranes. (**A**) Transmission electron micrograph of highpressure frozen, resin-embedded, thin section of a cell periphery showing a plastid (P) profile within the cytoplasm (Cy). (**B**) Cryo-electron tomogram showing the membranes separating the plastid thylakoids (Th, black arrowhead) and stroma from the cytoplasm. White dashed boxed region is magnified in b) where white arrowheads indicate four bounding membranes of the plastid. (**C**) Z-series of 5 successive virtual sections of the tomogram area shown by the dashed box in (B). The bounding membranes show the outer and inner membrane pairs maintaining an approximately fixed separation distance between each membrane, but variation in the spacing between these pairs is seen (asterisk). Nu, nucleus; M, mitochondrion; Cy, cytoplasm; P, plastid; PM, plasma membrane; Th and black arrowheads, thylakoids.

### The ancestral dinoflagellate peridinin plastid lacks SELMA and this plastid likely persists in Kareniaceae with haptophyte plastids

The ancestral photosynthetic plastid of dinoflagellates contains a distinctive secondary pigment, the carotenoid peridinin ^23,24^. This ‘peridinin plastid’ is surrounded by only three membranes which distinguishes it from the four membrane-bound plastids of most other organisms with complex plastids ^25^. In none of our phylogenetic analyses were SELMA orthologues found in dinoflagellates that only contain this ancestral plastid, with 45 species sampled across 30 genera (Figs 1-3, S1). The ERAD paralogues of the ER, on the other hand, were ubiquitously present in these taxa showing that available sequence coverage was not limiting these searches. Furthermore, our analyses show that in the related apicomplexans even the basal groups with a plastid retain SELMA proteins (e.g., the marosporidian *Rhytidocystis*, gregarines *Selendinium* and *Siedleckia*, chromopodellid *Piridium*, and squirmid *Digyalum*) whereas taxa that have lost the plastid also lack SELMA orthologues (*Cryptosporidium* spp. and gregarines *Cephaloidophora* and *Heliospora*) (Figs 2&3, S1). In the dinoflagellates with the peridinin plastid, orthologues of Tic110, Tic20 and Tic22 were consistently found, indicating that this plastid likely only requires the canonical TOC/TIC for protein translocation across the inner membranes. Vesicular fusion from the endomembrane system is known to deliver proteins across the outermost membrane (**Figure 6**) ^26^. Whether SELMA occurred previously in dinoflagellates but was lost along with the third membrane, or was never present in these cells, is unknown.

**Figure 6:**
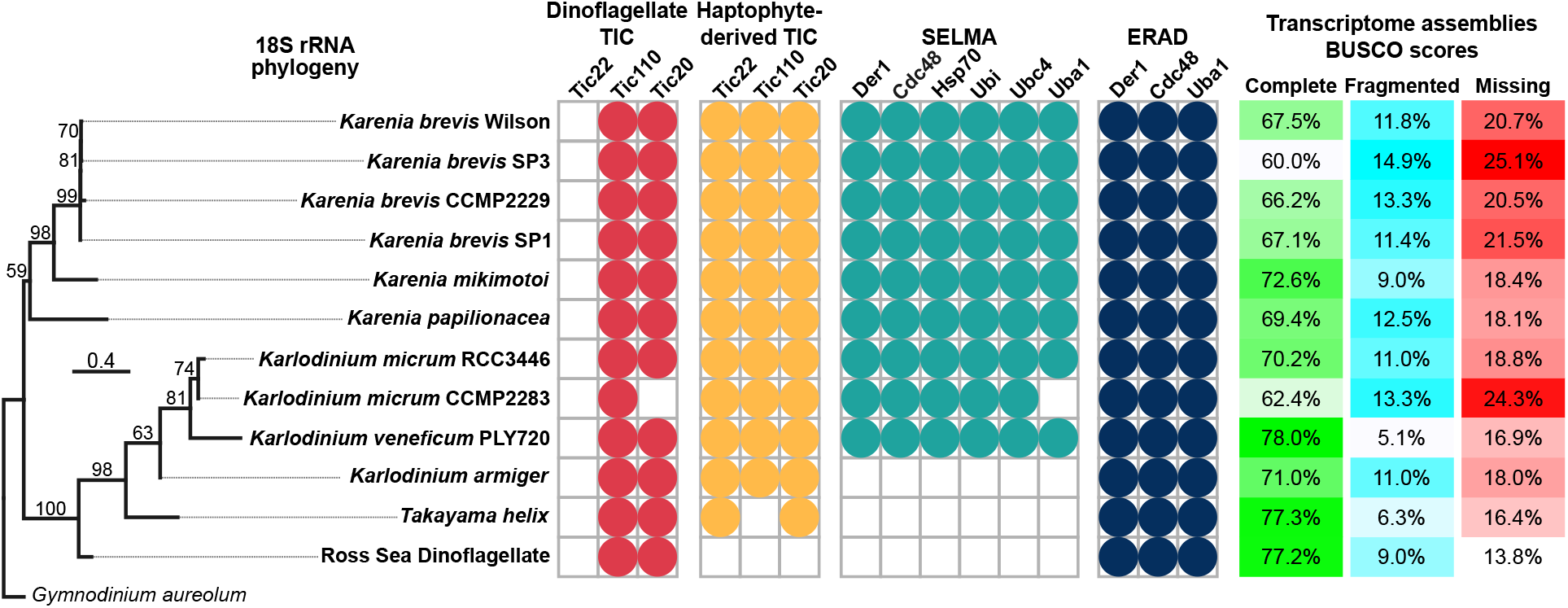
Presence and absence of plastid translocons in Kareniaceae. Phylogeny of Kareniaceae dinoflagellates and the detection of expressed plastid translocon proteins in each taxon indicated by colored circles. BUSCO scores as an estimate of transcriptome coverage are given for each.

Unexpectedly in Kareniaceae, peridinin-plastid orthologues of Tic110 and Tic20 were found in addition to orthologues derived from haptophytes (**Figures 1B&C, 6**). These data suggest the presence of two phylogenetically distinct plastids in Kareniaceae where a cryptic relic of the peridinin plastid has been retained despite the presence of the new haptophyte-derived plastids for photosynthesis.

### SELMA loss has occurred in instances of further plastid replacement in Kareniaceae

In two Kareniaceae species, further instances of plastid gain have occurred in apparent like-for-like plastid replacements. Phylogenies of plastid-encoded genes show that *Karlodinium armiger* and *Takayama helix* have plastids more closely related to *Prymnesium* and *Phaeocystis* haptophytes, respectively, than those of the other *Karlodinium* or *Karenia* spp. ^9^. In both cases, previously acquired nucleus-encoded haptophyte-derived genes apparently continue to service these new plastids ^9^. This indicates that many or all nucleus-encoded plastid genes were compatible with the replacement plastids when taken from closely related groups (i.e., other haptophytes), and that these genes provide a preadaptation for possible ongoing plastid replacements. Indeed, we see in *K. armiger* that even the TICs for protein import are of common origin with the other *Karlodinium* spp. and, therefore, have also facilitated plastid replacement (**Figure S1**). Surprisingly, both *K. armiger* and *T. helix* lack any genes for SELMA (**Figures 6, S1**). These losses of the SELMA complex resemble the peridinin plastid that is surrounded by only three membranes. This suggests that the acquisition of new plastids in both *K. armiger* and *T. helix* might have involved the elimination of one bounding membrane. Bipartite plastid-targeting sequences are used for both three- and four-membrane plastids ^19^. Therefore, the loss of the equivalent of the third (SELMA) membrane would not be predicted to disrupt protein import of existing nucleus-encoded proteins.

A further Kareniaceae species, the Ross Sea Dinoflagellate lacks a stable haptophyte-derived plastid but does acquire and maintain temporary haptophyte endosymbionts captured from *Phaeocystis* spp. ^5^. While some haptophyte genes occur in the nucleus of this dinoflagellate ^6^, we detected no expression of haptophyte SELMA or even TIC proteins in their transcriptomes (**Figure 6**). Thus, any protein import in this system must rely on the translocons acquired in situ with the ‘kleptoplast’, and they would seemingly not be able to be resynthesized if damaged or degraded

### Dinoflagellates with additional paralogous ERAD machineries

A different group of dinoflagellates (family Kryptoperidiniaceae of the Peridiniales) have also acquired new alternative plastids, in this case derived from diatom endosymbionts. These so-called ‘dinotom’ dinoflagellates stably maintain, replicate and inherit these new endosymbionts ^24^. Dinotoms are unusual, however, in that a further single membrane is present surrounding their endosymbionts creating a fifth membrane separating the plastid lumen from host cytosol ^27^. No protein targeting from the host cell to endosymbiont has apparently been established ^28^. Thus, these endosymbionts maintain a diatom nucleus that encodes and expresses most plastid-related proteins that then translocate across the existing four diatom membranes surrounding the stroma and thylakoids. Given the relative intactness of the diatom as endosymbiont, we hypothesised that the diatom SELMA machinery would have been retained to enable protein targeting of the diatom-encoded plastid proteins. Our phylogenies of all the SELMA proteins, as well as the TIC proteins, showed that stramenopile-grouping orthologues for plastid import are all present in the dinotom *Kryptoperidinium triquetrum*, consistent with maintenance of the diatom-derived machinery (Figs 1-3, S1). *K. triquetrum* also possesses the ERAD proteins that group with the dinoflagellate ERAD orthologues as expected for the presence of the canonical ER machinery. However, an additional set of ERAD paralogues is also present in *K. triquetrum*, and these proteins group with the diatom/stramenopile ERAD proteins (Figs 1-3, S1). These data indicate that the ER of the diatom symbiont still retains its ERAD protein quality control processes. Thus, dinotoms maintain three paralogous ERAD machineries—two serving two functional but separate ER systems, and one (SELMA) representing the repurposed plastid translocon— and provides a second example of ERAD duplication through endosymbiosis.

### Kareniaceae present a model for serial red plastid acquisition and exchange

This study shows that plastids in the Kareniaceae dinoflagellates have not accrued further membranes during their acquisition from haptophytes, and that these plastids maintained all the necessary translocons from the source haptophytes for protein import (**Figure 7**). Effectively, these plastid organelles occur and function in the cytosol of Kareniaceae dinoflagellates as they would have previously in the cytosol of haptophytes. Most genes for haptophyte-derived plastid proteins are known to have been transferred to the dinoflagellate nuclei ^9,28–31^ and, given that they would have possessed pre-existing bipartite leaders for targeting to haptophyte plastids, their expression in the new host could facilitate protein uptake into this plastid as before. The sorting of plastid proteins within the ER/endomembrane system to target them to the plastid would be the only necessary adaptation required after the acquisition of these new plastids, and it is hard to predict how difficult this might be to develop. But in the Kareniaceae, this process might have also exploited existing protein sorting routes used for proteins of the relict peridinin plastid that our data suggests still cooccurs today. The discrimination of proteins required in the two different plastids might have been assisted through the SELMA-based selection of proteins for the haptophyte plastid that is absent for the peridinin plastid. In any case, it is apparent that little or no new translocation machinery needed to be developed for a pre-existing complex red-type plastid to be transferred to a different eukaryotic lineage.

**Figure 7.**
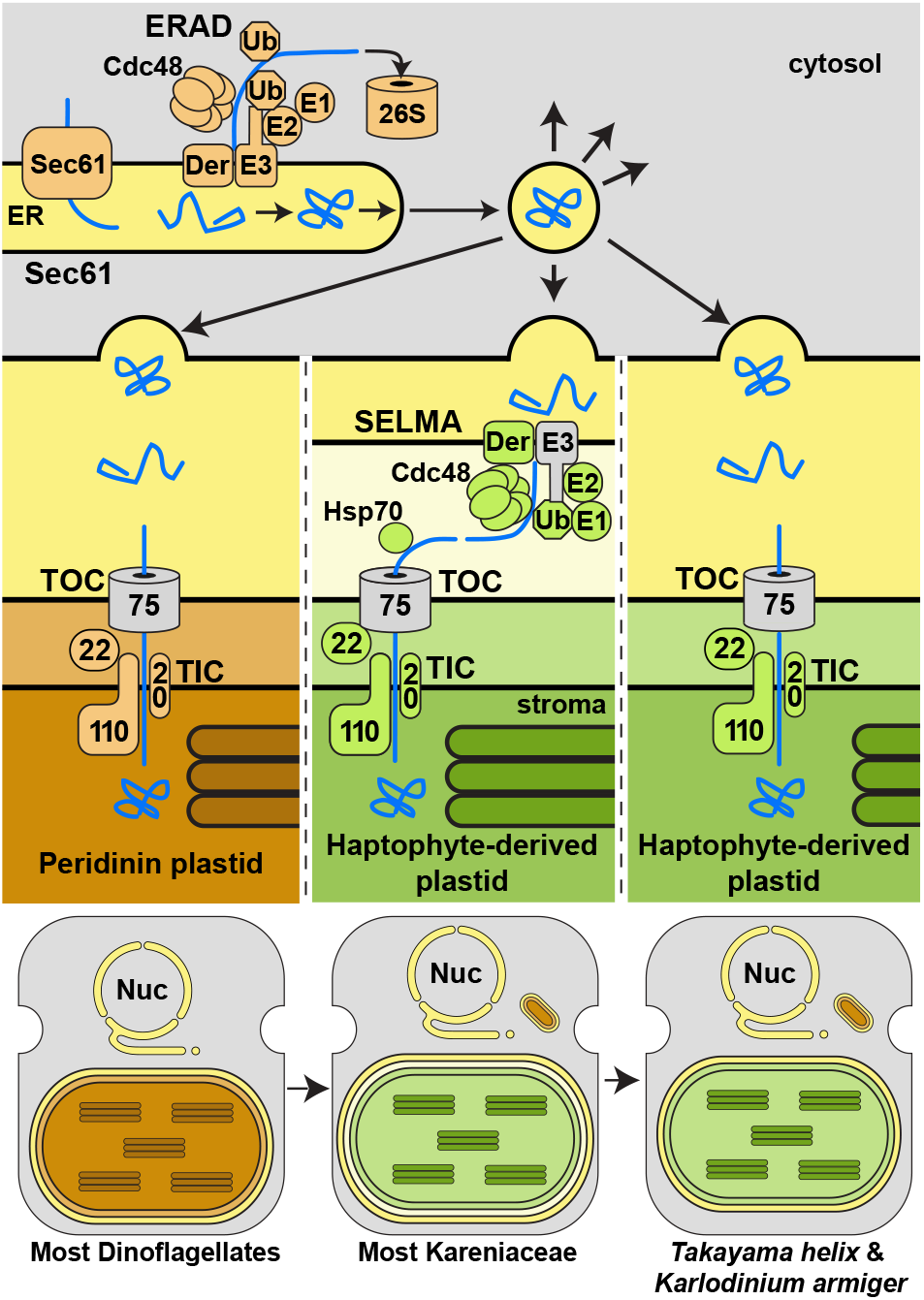
Model of ancestry and evolution of plastid protein translocation machinery in Kareniaceae dinoflagellates. Ancestral peridinin plastids of dinoflagellates are surrounded by three membranes and protein targeting occurs via the ER translocon Sec61 and vesicle delivery. Upon outer membrane fusion, protein cargo then passes through the TOC and TIC complexes. Most known Kareniaceae dinoflagellates inherited new plastids derived from haptophytes, and likely adopted the dinoflagellate routing through the ER to the outermost plastid membrane. From here, haptophyte-derived translocons (green), including SELMA, complete protein import. Further plastid replacement in *T. helix* and *K. armiger* with alterative haptophyte plastids have maintained the preexisting TICs but eliminated all SELMA components, and presumably the equivalent plastid membrane. Grey translocons remain unidentified.

This model for plastid exchange presented by the Kareniaceae suggests that an intact haptophyte algal endosymbiont likely never occurred as an intermediate stage in their evolution, unlike that seen for the seemingly stalled diatom endosymbionts of Kryptoperidiniaceae dinotoms. In dinotoms the additional (fifth) membrane around the diatom symbiont might present one protein trafficking challenge too many for the development of a protein import process ^32^. The persistence of diatom nuclei, ERAD and SELMA indicate that dinotom plastid proteins still co-translationally enter the diatom ER *en route* to the plastid. This process is likely incompatible with any further preceding trafficking events across the outermost fifth membrane and from the dinoflagellate cytosol. A consequence of the model of direct organelle procurement inferred for Kareniaceae is that the transfer of plastid genes to the nucleus of the new host would have to occur by way of repeated feeding events on the same or closely related algal taxa rather than from a constantly maintained symbiont nucleus. The Ross Sea Dinoflagellate, as well as the euglenid *Rapaza*, provide examples of this process where temporary kleptoplasts receive some maintenance from small numbers of genes acquired in the host’s nucleus by horizontal gene transfer from their prey’s nuclei ^6,33^. This process has seemingly continued to completion in the Kareniaceae where the once kleptoplasts are now permanent stable organelles with no persistent haptophyte nuclei.

A corollary of the protracted process of endosymbiont gene gain is that a long history of constant organelle gain and turnover would have occurred. Indeed, there is evidence that different *Karlodinium* and *Karenia* spp. containing similar, but taxonomically distinct, haptophyte plastids, suggesting that they ultimately fixed different stable endosymbionts ^5,9^. *K. armiger* and *T. helix*, provides further evidence of the ongoing transmissibility of complex red plastids, each possessing a more recently replaced version of a haptophyte-derived plastid. Kareniaceae dinoflagellates remain eukaryovorous, sucking up prey organelles by myzocytotic feeding, and it is plausible that further cases of plastid replacement will be identified in this group ^34^. It is curious that *K. armiger* and *T. helix* both lost SELMA with their plastid replacements, and presumably the third membrane which SELMA is required to translocate proteins across. This apparent reversion to the state of the peridinin plastid raises the question of whether the canonical dinoflagellate three-membrane bound plastid is an equivalent replacement of a possible ancestral SELMA-containing four-membrane plastid as is found throughout the sister lineage, Apicomplexa.

In principle, we see no reason why the model of complex red plastid gain in the Kareniaceae might not also account for the gains of equivalent plastids in haptophytes, stramenopiles and apicomplexans, and perhaps even separate gains in chrompodellids *Chromera* and *Vitrella*. In all cases the same number of membranes surround the plastid, and the same translocons and trafficking steps occur in all ^14,19,22,35^. The possible number of exchanges of serial secondary plastids, and their directions of travel, might be difficult to gauge. But this model provides a mechanistic solution to the ‘Rhodoplex’ hypothesis ^36^ that posits the gain of much of aquatic photosynthetic diversity through multiple exchanges of a single red algal-derived secondary plastid ^3,37–41^ as an alternative to the now phylogenetically impossible single ancestral gain predicted by the original Chromalveolate hypothesis.

## ACKNOWLEDGEMENTS

We thank Elisabeth Hehenberger for providing assembled transcriptome datasets for the Ross Sea Dinoflagellate, Katherine Helliwell and Angela Ward for providing *Karlodinium veneficum* PLY720 cultures and advice on their growth, and Dima Chirgadze and Lee Cooper for assistance with cryo-electron tomography. This work was supported by grants from the Gordon and Betty Moore Foundation to RFW (doi:10.37807/GBMF9194) and JD (doi:10.37807/GBMF11532). GB was supported by a Herchel Smith Postdoctoral Research Fellowship.

## AUTHOR CONTRIBUTIONS

RFW conceived the study with contributions by WHL and LK. WHL performed cell culturing, RNA extraction, transcriptome assembly and the phylogenetic analyses with LK and SL undertaking preliminary analyses. WHL analysed protein targeting signals with contributions by LK. VF contributed to the data analysis strategy and managed computational resources. GP performed cryoET imaging, data processing and tomogram analysis under the supervision of BFL. GB processed cell samples for FIB-milling, and TD performed the FIB-milling and provided advice on tomography. BG, DY and JD generated the TEM data. RFW and WHL wrote the manuscript with input from all authors.

## DECLARATION OF INTERESTS

The authors declare no competing interests.

## METHODS

### Cultivation of *Karlodinium veneficum* PLY720

*Karlodinium veneficum* PLY720 was obtained from The Plymouth Culture Collection of Marine Algae, Marine Biological Association, Plymouth, UK. The culture collection records report that this strain was originally isolated in 1976 from a marine sample collected from a fjord in Norway (59°30’N 10°36’E), and that it is synonymous with the strains CCMP415 held at the National Center for Marine Algae and Microbiota, Bigelow, USA, and NEPCC 734 held at The Canadian Center for the Culture of Microorganisms, Vancouver, Canada. Cells were grown in L1 medium, typically in 200 ml volumes in T175 culture flasks. Culture flasks were maintained in an incubator with a 14:10 hours light/dark cycle, with a light intensity of approximately 20 µmol m^-2^ s^-1^ provided by LEDs, and at a consistent temperature of 20 ^°^C and sub-cultured monthly by inoculating filter-sterilised L1 medium with mature culture, typically in a volume ratio of 40:1.

### RNA extraction

To prepare samples for long-read sequencing cells were harvested from two 200 ml *Karlodinium veneficum* PLY720 cultures were harvested; one six hours into the light phase and one four hours into the dark phase of a culture light/dark cycle. Samples for short-read sequencing were prepared as part of a concurrent unpublished study in which cells from twenty-one 200 ml *Karlodinium veneficum* PLY720 cultures were harvested. In all cases cells from 200 ml cultures were collected by decanting the cultures into four 50 ml tubes, which were then centrifuged at 5000 x g for 5 minutes at 4 ° C. The supernatants were then discarded, and the pellets pooled by transferring to a 2 ml microcentrifuge tube. This tube was then centrifuged using the same conditions as the first centrifugation, after which the supernatant was discarded and the tube containing the remaining cell pellet was flash-frozen in liquid N2 and then stored at -80 ^°^C until further processing. All samples for both long and short read sequencing were then processed in the same way to extract RNA using TRIzol, with a detailed step-by-step description of the methods deposited at protocols.io (dx.doi.org/10.17504/protocols.io.n92ld9xe7g5b/v1).

### Library preparation, sequencing, transcriptome assembly and gene prediction

PacBio long-read sequencing was performed by the Earlham Institute, Norwich, UK. This included the preparation of sequencing libraries from the extracted RNA using the PacBio Iso-Seq Express Template Preparation (v2, no size-selection) kit, sequencing the libraries using a PacBio Sequel II SMRT cell (8M, v2, 30hr Movie), and performing IsoSeq3 analysis to process and refine the raw reads and generate HiFi reads. Illumina short read sequencing was performed by the Genomics department at Cancer Research UK Cambridge Institute. This included preparation of sequencing libraries using the Illumina Stranded mRNA Prep Kit, which were then sequenced using an Illumina NovaSeq 6000 system with a SP flowcell to generate 2 × 50 bp (paired-end) reads. Long reads and short reads were co-assembled using Trinity version 2.11.0 ^42^ and gene prediction performed using TransDecoder version 5.5.0 (https://github.com/TransDecoder/transdecoder.github.io.git).

### BUSCO analysis

BUSCO analysis was performed on predicted peptide files for all twelve Kareniaceae transcriptomes investigated in the present study using Busco version 5.5.0 ^43^ with the eukaryota_odb10 busco reference dataset.

### Database sampling and phylogenetic analysis

A detailed description of the methodology and workflow, including the code used to perform the phylogenetic analyses for translocon proteins is deposited at GitHub (https://github.com/camwallerlab/Methods-forphylogenetic-analysis-of-plastid-translocons). Briefly, this involved creating a custom protein sequence database from the datasets listed in Supplementary Table S1 obtained from UniProt, the MMETSP reassemblies (https://doi.org/10.5281/zenodo.3247846) ^44^, and VEuPathDB ^45^, as well as transcriptomes for organisms sequenced in three previous studies ^9,46,47^ and the *Karlodinium veneficum* PLY720 transcriptome generated in the present study. This custom database was then searched using blastp (BLAST+ version 2.11.0) for homologues of each of the translocon proteins investigated in the present study using characterised versions of these proteins as queries. The homologues obtained from these searches were then clustered with cdhit (CD-HIT version 4.8.1) ^48^ to remove highly similar sequences, thereby reducing overall redundancy of the dataset. Iterative rounds of alignment using mafft (MAFFT version 7.475) ^49^, conserved site selection using trimal (trimAl version 1.4) ^50^, tree inference with FastTreeMP (FastTree version 2.1.11) ^51^ using the default settings and manual removal of sequences were then performed. The purpose of these iterations was to further reduce dataset redundancy, as well as to identify and remove dissimilar or poorly aligning sequences. The final curated datasets for each translocon protein that these methods obtained were then aligned using mafft-linsi (MAFFT version 7.475) and conserved sites were selected using trimal (trimAl version 1.4). A phylogeny was then inferred for each translocon dataset using the program iqtree2 (IQ-TREE version 2.1.2) ^51^ with 1000 ultrafast bootstrap replicates (UFBoot2) ^52^ and using the best-fitting model, LG+F+I+G4 ^53,54^that was chosen according to the Bayesian Information Criterion by ModelFinder ^55^, all implemented within iqtree2.

For the 18S rRNA gene phylogeny, nucleotide sequences for each of the Kareniaceae species studied were extracted from the corresponding transcriptome assemblies using barrnap (Barrnap version 0.9, https://github.com/tseemann/barrnap). Sequences were aligned using mafft-linsi (MAFFT version 7.475) and conserved sites were selected using trimal (trimAl version 1.4). A tree was then inferred from the trimmed alignment using the program iqtree2 (IQ-TREE version 2.1.2) with 1000 ultrafast bootstrap replicates and using the best-fitting model, TN+F+I ^56^, that was chosen according to the Bayesian Information Criterion by ModelFinder, all implemented within iqtree2.

### Sample preparation for transmission electron microscopy

High-pressure freezing (HPM100, Leica) followed by freeze substitution (EM ASF2, Leica) was conducted to prepare samples for electron microscopy following the published protocols ^57^. Cultured *Karlodinium micrum* PLY720 cells were harvested at exponential growth phase and concentrated by 2 min centrifugation at 2,500 g prior to cryo-fixation with high pressure freezing. Resin blocks were obtained after the freeze substitution ^57^. For TEM analysis, ultrathin sections of 60 nm thickness were mounted onto copper grids or slots coated with formvar and carbon. Sections were then stained in 1% uranyl acetate (10 min) and lead citrate (5 min). Micrographs were obtained using a Tecnai G2 Spirit BioTwin microscope (FEI) operating at 120 kV with an Orius SC1000 CCD camera (Gatan).

### Cryo-FIB lamella preparation, cryo-ET data collection and processing

*Karlodinium micrum* PLY720 cells were allowed to settle in a 10 μl aliquot upon a poly-L-lysine coated holey carbon film gold grid that was previously glow-discharged for 60 s. Grids were sealed in a plastic container and transferred to an incubator and after 2 hrs, a second aliquot was added. Before vitrification, grids were manually blotted on the reverse side for 2 s, then plunged into liquid ethane with an FEI Vitrobot (100% humidity) and stored in liquid nitrogen until used. Grids were first screened in a Talos Arctica operated at 200 kV (equipped with a Falcon 3EC detector) and grids of good quality were chosen for FIB-milling. For milling, the grid was transferred to a Zeiss crossbeam 550 Gemini 2 system and a layer of platinum was sputtered to the surface of sample for 1 min 20 s. Rough milling was done automatically at current range of 700-100 pA, and polishing was done manually at 50 pA, aiming at a lamellae thickness of 180nm. An additional Pt layer was added to stabilise lamella for 2 seconds at 5 mA.

The thin lamellae were imaged on a Titan Krios G2 transmission electron microscope (Thermo Fisher Scientific/FEI) operated at 300 kV equipped with a Gatan K3 direct electron detector. Tilt series were acquired with a dose-symmetric scheme, with a 3º increment between +60º and -60º. A pixel size of 3.5 Å/pixel was used, with a defocus range from -3.0 to -5.0. Images were acquired with a total dose of 73.4 e^−^ (Å) ^−2^. Gain-correction, motion correction and defocus estimation were performed in WARP 1.0.9 ^58^. The tilt series were aligned in AreTomo ^59^, and tomograms were reconstructed in AreTomo with a binning of 8 (28 Å/pixel) and using simultaneous algebraic reconstruction technique (SART) ^60^.

## Data Availability

Raw sequencing data used to assemble the *Karlodinium veneficum* PLY720 reference transcriptome is available for download from the NCBI Sequence Read Archive under the BioProject accession PRJNA1117636 (BioSample accession SAMN41577327). Assembled transcriptome data, including transcripts and predicted proteins in fasta format, in addition to files that were used to generate phylogenies, including unaligned sequences, aligned sequences, trimmed alignments, and output files from IQTREE 2, are available at Figshare (https://figshare.com/s/41c2e3c38d039359816c, DOI: 10.6084/m9.figshare.21602697)

## Notes

### Competing Interest Statement

The authors have declared no competing interest.

## REFERENCES

1. Cavalier-Smith, T. (1999). Principles of protein and lipid targeting in secondary symbiogenesis: euglenoid, dinoflagellate, and sporozoan plastid origins and the eukaryote family tree. J. Eukaryot. Microbiol. 46, 347–366. 10.1111/j.1550-7408.1999.tb04614.x.

2. Burki, F., Roger, A.J., Brown, M.W., and Simpson, A.G.B. (2019). The new tree of eukaryotes. Trends Ecol & Evol, 1–13. 10.1016/j.tree.2019.08.008.

3. Strassert, J.F.H., Irisarri, I., Williams, T.A., and Burki, F. (2021). A molecular timescale for eukaryote evolution with implications for the origin of red algal-derived plastids. Nat Comms 12, 1879. 10.1038/s41467-021-22044-z.

4. Takahashi, K., Benico, G., Lum, W.M., and Iwataki, M. (2019). Gertia stigmatica gen. et sp. nov. (Kareniaceae, Dinophyceae), a new marine unarmored dinoflagellate possessing the peridinin-type chloroplast with an eyespot. Protist 170, 125680. 10.1016/j.protis.2019.125680.

5. Gast, R.J., Moran, D.M., Dennett, M.R., and Caron, D.A. (2007). Kleptoplasty in an Antarctic dinoflagellate: caught in evolutionary transition? Environ Microbiol 9, 39–45. 10.1111/j.1462-2920.2006.01109.x.

6. Hehenberger, E., Gast, R.J., and Keeling, P.J. (2019). A kleptoplastidic dinoflagellate and the tipping point between transient and fully integrated plastid endosymbiosis. PNAS 116, 17934–17942. 10.1073/pnas.1910121116.

7. Yoon, H.S., Hackett, J.D., and Bhattacharya, D. (2002). A single origin of the peridinin- and fucoxanthin-containing plastids in dinoflagellates through tertiary endosymbiosis. PNAS 99, 11724–11729. 10.1073/pnas.172234799.

8. Tengs, T., Dahlberg, O.J., Shalchian-Tabrizi, K., Klaveness, D., Rudi, K., Delwiche, C.F., and Jakobsen, K.S. (2000). Phylogenetic analyses indicate that the 19’Hexanoyloxy-fucoxanthin-containing dinoflagellates have tertiary plastids of haptophyte origin. Mol Biol Evol 17, 718–729.

9. Vanclová, A.M.N., Nef, C., Füssy, Z., Vancl, A., Liu, F., Bowler, C., and Dorrell, R.G. (2024). New plastids, old proteins: repeated endosymbiotic acquisitions in kareniacean dinoflagellates. EMBO Rep. 25, 1859–1885. 10.1038/s44319-024-00103-y.

10. Bergholtz, T., Daugbjerg, N., Moestrup, Ø., and Fernández-Tejedor, M. (2006). On the identity of Karlodinium veneficum and description of Karlodinium armiger sp. nov. (dinophyceae), based on light and electron microscopy, nuclear-encoded LSU rDNA, and pigment composition1. J Phycol 42, 170–193. 10.1111/j.1529-8817.2006.00172.x.

11. Stork, S., Lau, J., Moog, D., and Maier, U.-G. (2013). Three old and one new: protein import into red algal-derived plastids surrounded by four membranes. Protoplasma 250, 1013–1023. 10.1007/s00709-013-0498-7.

12. Jin, Z., Wan, L., Zhang, Y., Li, X., Cao, Y., Liu, H., Fan, S., Cao, D., Wang, Z., Li, X., et al. (2022). Structure of a TOC-TIC supercomplex spanning two chloroplast envelope membranes. Cell 185, 4788-4800.e13. 10.1016/j.cell.2022.10.030.

13. Glaser, S., Dooren, G.G. van, Agrawal, S., Brooks, C.F., McFadden, G.I., Striepen, B., and Higgins, M.K. (2012). Tic22 is an essential chaperone required for protein import into the apicoplast. J. Biol. Chem. 287, 39505–39512. 10.1074/jbc.m112.405100.

14. Stork, S., Moog, D., Przyborski, J.M., Wilhelmi, I., Zauner, S., and Maier, U.-G. (2012). Distribution of the SELMA translocon in secondary plastids of red algal origin and predicted uncoupling of ubiquitin-dependent translocation from degradation. Euk Cell 11, 1472–1481. 10.1128/ec.00183-12.

15. Agrawal, S., Chung, D.-W.D., Ponts, N., Dooren, G.G. van, Prudhomme, J., Brooks, C.F., Rodrigues, E.M., Tan, J.C., Ferdig, M.T., Striepen, B., et al. (2013). An apicoplast localized ubiquitylation system is required for the import of nuclear-encoded plastid proteins. PLOS Paths 9, e1003426. 10.1371/journal.ppat.1003426.

16. Fellows, J.D., Cipriano, M.J., Agrawal, S., and Striepen, B. (2017). A plastid protein that evolved from ubiquitin and is required for apicoplast protein import in Toxoplasma gondii. mBio 8. 10.1128/mbio.00950-17.

17. Sommer, M.S., Gould, S.B., Lehmann, P., Gruber, A., Przyborski, J.M., and Maier, U.-G. (2007). Der1-mediated preprotein import into the periplastid compartment of chromalveolates? Mol Biol Evol 24, 918–928. 10.1093/molbev/msm008.

18. Felsner, G., Sommer, M.S., Gruenheit, N., Hempel, F., Moog, D., Zauner, S., Martin, W., and Maier, U.G. (2011). ERAD components in organisms with complex red plastids suggest recruitment of a preexisting protein transport pathway for the periplastid membrane. Genome Biol Evol 3, 140–150. 10.1093/gbe/evq074.

19. Patron, N.J., and Waller, R.F. (2007). Transit peptide diversity and divergence: A global analysis of plastid targeting signals. BioEssays 29, 1048–1058. 10.1002/bies.20638.

20. Käll, L., Krogh, A., and Sonnhammer, E.L.L. (2004). A combined transmembrane topology and signal peptide prediction method. J Mol Biol 338, 1027–1036. 10.1016/j.jmb.2004.03.016.

21. Crooks, G.E., Hon, G., Chandonia, J.-M., and Brenner, S.E. (2004). WebLogo: A sequence logo generator. Genome Res. 14, 1188–1190. 10.1101/gr.849004.

22. Gould, S.B., Maier, U.-G., and Martin, W.F. (2015). Protein import and the origin of red complex plastids. Curr Biol 25, R515–21. 10.1016/j.cub.2015.04.033.

23. Rapoport, H., Strain, H.H., Svec, W.A., Aitzetmueller, K., Grandolfo, M., Katz, J.J., Kjoesen, H., Norgard, S., and Liaaen-Jensen, S. (2002). Structure of peridinin, the characteristics dinoflagellate carotenoid. JACS 93, 1823–1825. 10.1021/ja00736a065.

24. Waller, R.F., and Koreny, L. (2017). Plastid complexity in dinoflagellates: a picture of gains, losses, replacements and revisions. In Secondary Endosymbioses. (Elsevier), pp. 105–143. 10.1016/bs.abr.2017.06.004.

25. Schnepf, E., and Elbrächter, M. (1999). Dinophyte chloroplasts and phylogeny-A review. Grana 38, 81–97.

26. Nassoury, N., Cappadocia, M., and Morse, D. (2003). Plastid ultrastructure defines the protein import pathway in dinoflagellates. J Cell Sci 116, 2867–2874. 10.1242/jcs.00517.

27. Tomas, R.N., and Cox, E.R. (1973). Observations on the symbiosis of Peridinium balticum and its intracellular alga. I. Ultrastructure. J Phycol 9, 304–323.

28. Burki, F., Imanian, B., Hehenberger, E., Hirakawa, Y., Maruyama, S., and Keeling, P.J. (2014). Endosymbiotic gene transfer in tertiary plastid-containing dinoflagellates. Euk Cell 13, 246–255. 10.1128/ec.00299-13.

29. Patron, N.J., Waller, R.F., and Keeling, P.J. (2006). A tertiary plastid uses genes from two endosymbionts. J Mol Biol 357, 1373–1382. 10.1016/j.jmb.2006.01.084.

30. Takishita, K., Ishida, K.-I., and Maruyama, T. (2004). Phylogeny of nuclear-encoded plastid-targeted GAPDH gene supports separate origins for the peridinin- and the fucoxanthin derivative-containing plastids of dinoflagellates. Protist 155, 447–458.

31. Ishida, K.-I., and Green, B.R. (2002). Second- and third-hand chloroplasts in dinoflagellates: phylogeny of oxygen-evolving enhancer 1 (PsbO) protein reveals replacement of a nuclear-encoded plastid gene by that of a haptophyte tertiary endosymbiont. PNAS 99, 9294–9299. 10.1073/pnas.142091799.

32. N, Y., WH, L., T, H., and RF, W. (2024). Dinotoms illuminate early pathways toward the stable acquisition of photosynthetic endosymbionts. In Endosymbiotic organelle acquisition, S. Schwartzbach, P. Kroth, and M, eds. (Springer, Cham).

33. Karnkowska, A., Yubuki, N., Maruyama, M., Yamaguchi, A., Kashiyama, Y., Suzaki, T., Keeling, P.J., Hampl, V., and Leander, B.S. (2023). Euglenozoan kleptoplasty illuminates the early evolution of photoendosymbiosis. PNAS 120, e2220100120. 10.1073/pnas.2220100120.

34. Yang, H., Hu, Z., Shang, L., Deng, Y., and Tang, Y.Z. (2020). A strain of the toxic dinoflagellate Karlodinium veneficum isolated from the East China Sea is an omnivorous phagotroph. Harmful Algae 93, 101775. 10.1016/j.hal.2020.101775.

35. Agrawal, S., Dooren, G.G. van, Beatty, W.L., and Striepen, B. (2009). Genetic evidence that an endosymbiont-derived endoplasmic reticulum-associated protein degradation (ERAD) system functions in import of apicoplast proteins. J Biol Chem 284, 33683–33691. 10.1074/jbc.m109.044024.

36. Petersen, J., Ludewig, A.-K., Michael, V., Bunk, B., Jarek, M., Baurain, D., and Brinkmann, H. (2014). Chromera velia, endosymbioses and the rhodoplex hypothesis--plastid evolution in cryptophytes, alveolates, stramenopiles, and haptophytes (CASH lineages). Genome Biol Evol 6, 666–684. 10.1093/gbe/evu043.

37. Bodył, A. (2018). Did some red alga-derived plastids evolve via kleptoplastidy? A hypothesis. Biol Rev 93, 201–222. 10.1111/brv.12340.

38. Bodył, A., Stiller, J.W., and Mackiewicz, P. (2009). Chromalveolate plastids: direct descent or multiple endosymbioses? Trends Ecol & Evol 24, 119-21-author reply 121–2. 10.1016/j.tree.2008.11.003.

39. Stiller, J.W. (2014). Toward an empirical framework for interpreting plastid evolution. J Phycol 50, 462–471. 10.1111/jpy.12178.

40. Ševčíková, T., Horák, A., Klimeš, V., Zbránková, V., Demir-Hilton, E., Sudek, S., Jenkins, J., Schmutz, J., Přibyl, P., Fousek, J., et al. (2015). Updating algal evolutionary relationships through plastid genome sequencing: did alveolate plastids emerge through endosymbiosis of an ochrophyte? Sci Rep 5, 10134. 10.1038/srep10134.

41. Stiller, J.W., Schreiber, J., Yue, J., Guo, H., Ding, Q., and Huang, J. (2014). The evolution of photosynthesis in chromist algae through serial endosymbioses. Nat Comms 5, 5764. 10.1038/ncomms6764.

42. Grabherr, M.G., Haas, B.J., Yassour, M., Levin, J.Z., Thompson, D.A., Amit, I., Adiconis, X., Fan, L., Raychowdhury, R., Zeng, Q., et al. (2011). Full-length transcriptome assembly from RNA-Seq data without a reference genome. Nat Biotech 29, 644–652. 10.1038/nbt.1883.

43. Manni, M., Berkeley, M.R., Seppey, M., Simão, F.A., and Zdobnov, E.M. (2021). BUSCO update: novel and streamlined workflows along with broader and deeper phylogenetic coverage for scoring of eukaryotic, prokaryotic, and viral genomes. Mol Biol Evol 38, 4647–4654. 10.1093/molbev/msab199.

44. Keeling, P.J., Burki, F., Wilcox, H.M., Allam, B., Allen, E.E., Amaral-Zettler, L.A., Armbrust, E.V., Archibald, J.M., Bharti, A.K., Bell, C.J., et al. (2014). The marine microbial eukaryote transcriptome sequencing project (MMETSP): illuminating the functional diversity of eukaryotic life in the oceans through transcriptome sequencing. PLOS Biol 12, e1001889. 10.1371/journal.pbio.1001889.

45. Alvarez-Jarreta, J., Amos, B., Aurrecoechea, C., Bah, S., Barba, M., Barreto, A., Basenko, E.Y., Belnap, R., Blevins, A., Böhme, U., et al. (2023). VEuPathDB: the eukaryotic pathogen, vector and host bioinformatics resource center in 2023. Nucl Acids Res 52, D808–D816. 10.1093/nar/gkad1003.

46. Mathur, V., Wakeman, K.C., and Keeling, P.J. (2021). Parallel functional reduction in the mitochondria of apicomplexan parasites. Curr Biol 31, 2920–2928.e4. 10.1016/j.cub.2021.04.028.

47. Janouškovec, J., Paskerova, G.G., Miroliubova, T.S., Mikhailov, K.V., Birley, T., Aleoshin, V.V., and Simdyanov, T.G. (2019). Apicomplexan-like parasites are polyphyletic and widely but selectively dependent on cryptic plastid organelles. eLife 8, 441. 10.7554/elife.49662.

48. Fu, L., Niu, B., Zhu, Z., Wu, S., and Li, W. (2012). CD-HIT: accelerated for clustering the next-generation sequencing data. Bioinformatics 28, 3150–3152. 10.1093/bioinformatics/bts565.

49. Katoh, K., and Standley, D.M. (2013). MAFFT multiple sequence alignment software version 7: improvements in performance and usability. Mol Biol Evol 30, 772–780. 10.1093/molbev/mst010.

50. Capella-Gutiérrez, S., Silla-Martínez, J.M., and Gabaldón, T. (2009). trimAl: a tool for automated alignment trimming in large-scale phylogenetic analyses. Bioinformatics 25, 1972–1973. 10.1093/bioinformatics/btp348.

51. Price, M.N., Dehal, P.S., and Arkin, A.P. (2010). FastTree 2 – Approximately maximum-likelihood trees for large alignments. PLoS ONE 5, e9490. 10.1371/journal.pone.0009490.

52. Hoang, D.T., Chernomor, O., Haeseler, A. von, Minh, B.Q., and Vinh, L.S. (2018). UFBoot2: improving the ultrafast bootstrap approximation. Mol Biol Evol 35, 518–522. 10.1093/molbev/msx281.

53. Le, S.Q., and Gascuel, O. (2008). An Improved general amino acid replacement matrix. Mol Biol Evol 25, 1307–1320. 10.1093/molbev/msn067.

54. Gu, X., Fu, Y.X., and Li, W.H. (1995). Maximum likelihood estimation of the heterogeneity of substitution rate among nucleotide sites. Mol Biol Evol 12, 546–557. 10.1093/oxfordjournals.molbev.a040235.

55. Kalyaanamoorthy, S., Minh, B.Q., Wong, T.K.F., Haeseler, A. von, and Jermiin, L.S. (2017). ModelFinder: fast model selection for accurate phylogenetic estimates. Nat Methods 14, 587–589. 10.1038/nmeth.4285.

56. Tamura, K., and Nei, M. (1993). Estimation of the number of nucleotide substitutions in the control region of mitochondrial DNA in humans and chimpanzees. Mol Biol Evol 10, 512–526. 10.1093/oxfordjournals.molbev.a040023.

57. Gallet, B., Moriscot, C., Schoehn, G., and Decelle, J. (2024). Cryo-fixation and resin embedding of biological samples for electron microscopy and chemical imaging. 10.17504/protocols.io.bp2l62kndgqe/v1.

58. Tegunov, D., and Cramer, P. (2019). Real-time cryo-electron microscopy data preprocessing with Warp. Nat Methods 16, 1146–1152. 10.1038/s41592-019-0580-y.

59. Zheng, S., Wolff, G., Greenan, G., Chen, Z., Faas, F.G.A., Bárcena, M., Koster, A.J., Cheng, Y., and Agard, D.A. (2022). AreTomo: An integrated software package for automated marker-free, motion-corrected cryo-electron tomographic alignment and reconstruction. J Struct Biol: X 6, 100068. 10.1016/j.yjsbx.2022.100068.

60. Andersen, A.H., and Kak, A.C. (1984). Simultaneous Algebraic Reconstruction Technique (SART): A superior implementation of the ART algorithm. Ultrason Imaging 6, 81–94. 10.1016/0161-7346(84)90008-7.

