## Supplementary figures and images for "Plastid translocon recycling in dinoflagellates demonstrates the portability of complex plastids between hosts"

### Figure S1

#### Cdc48

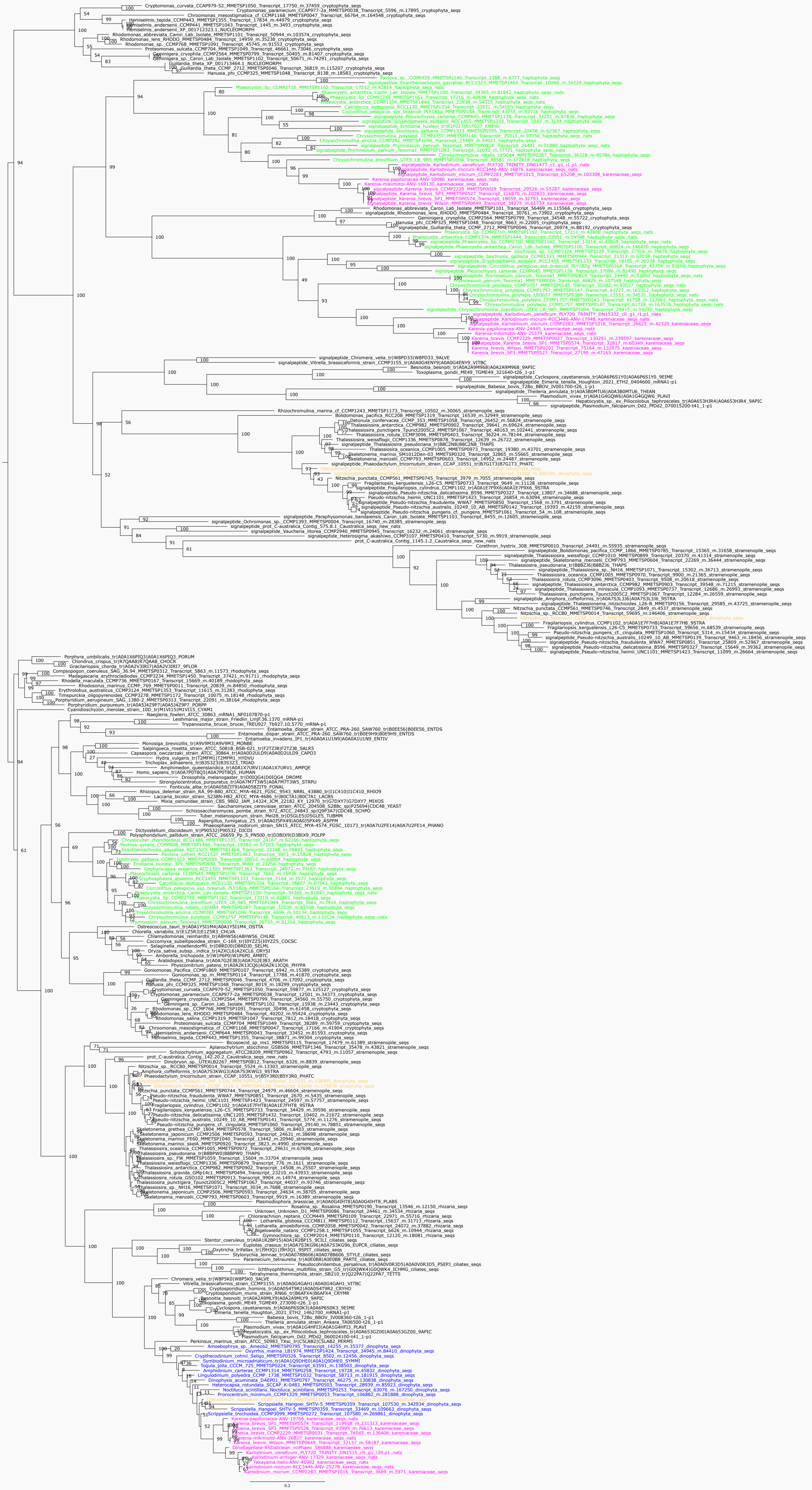

Der1

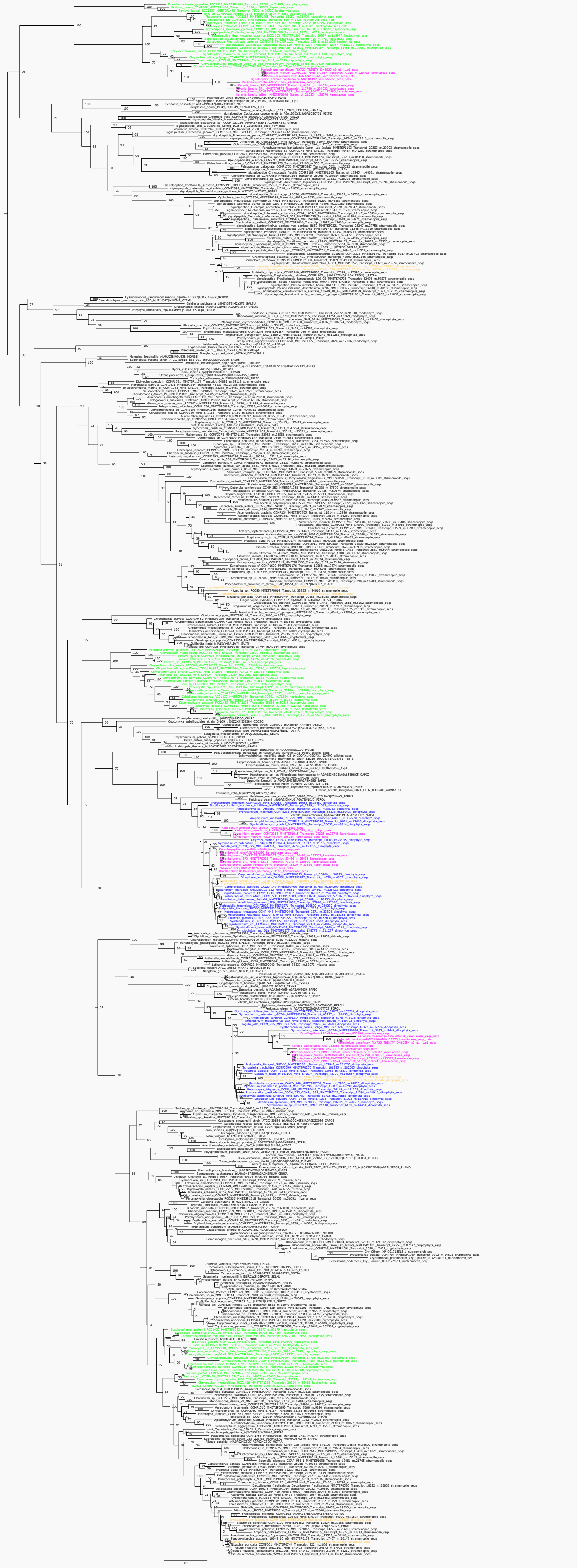

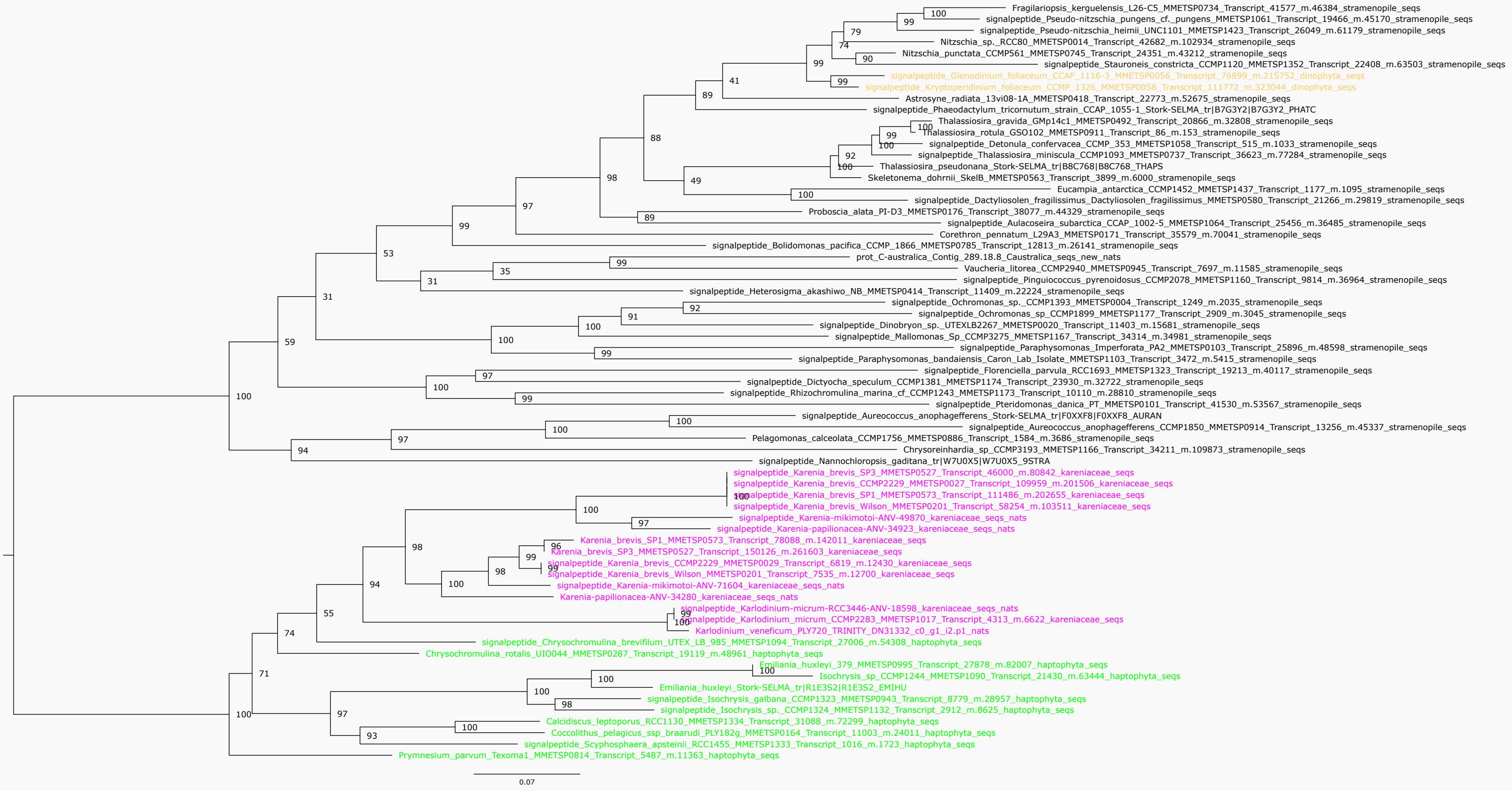

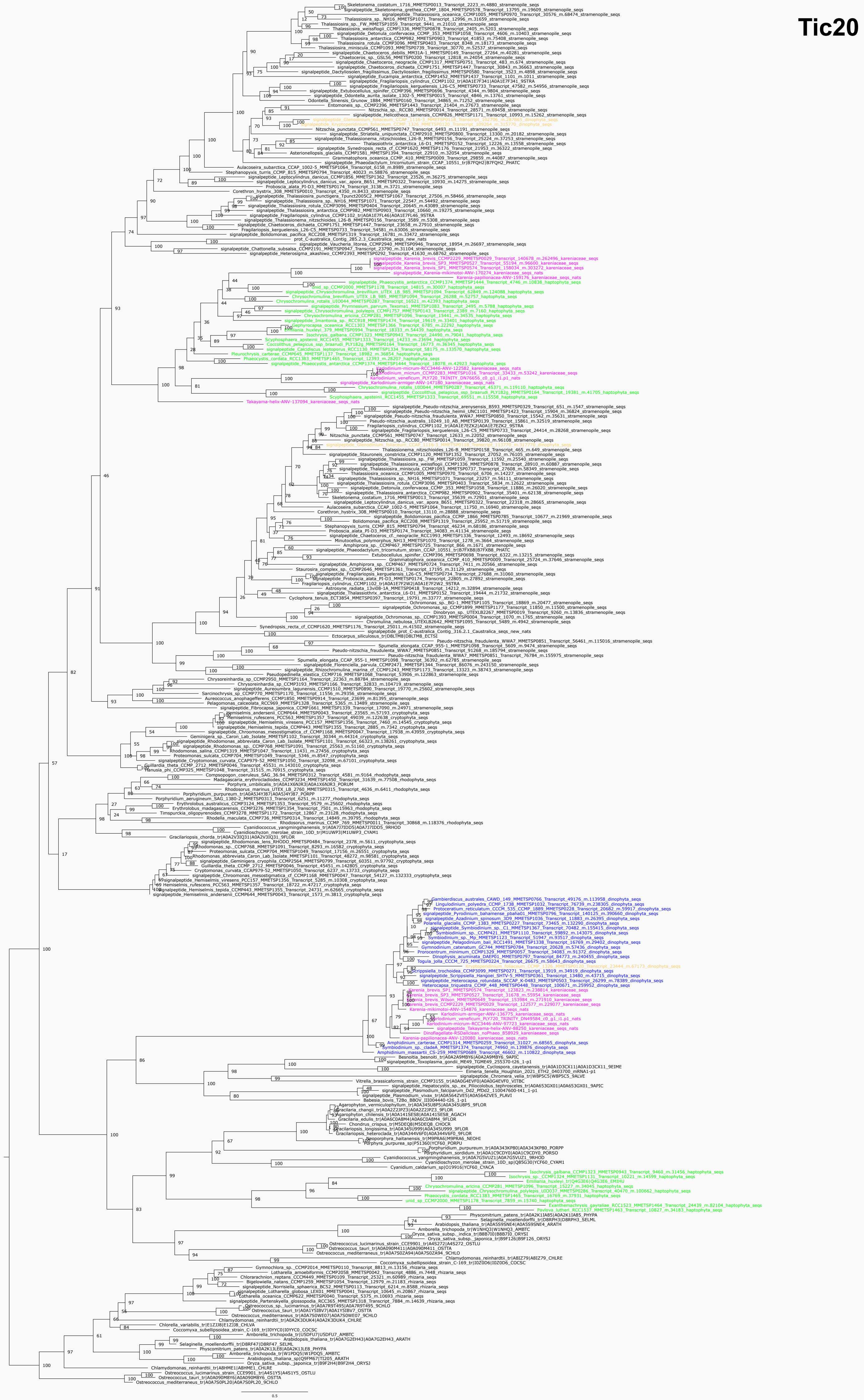

#### Tic22

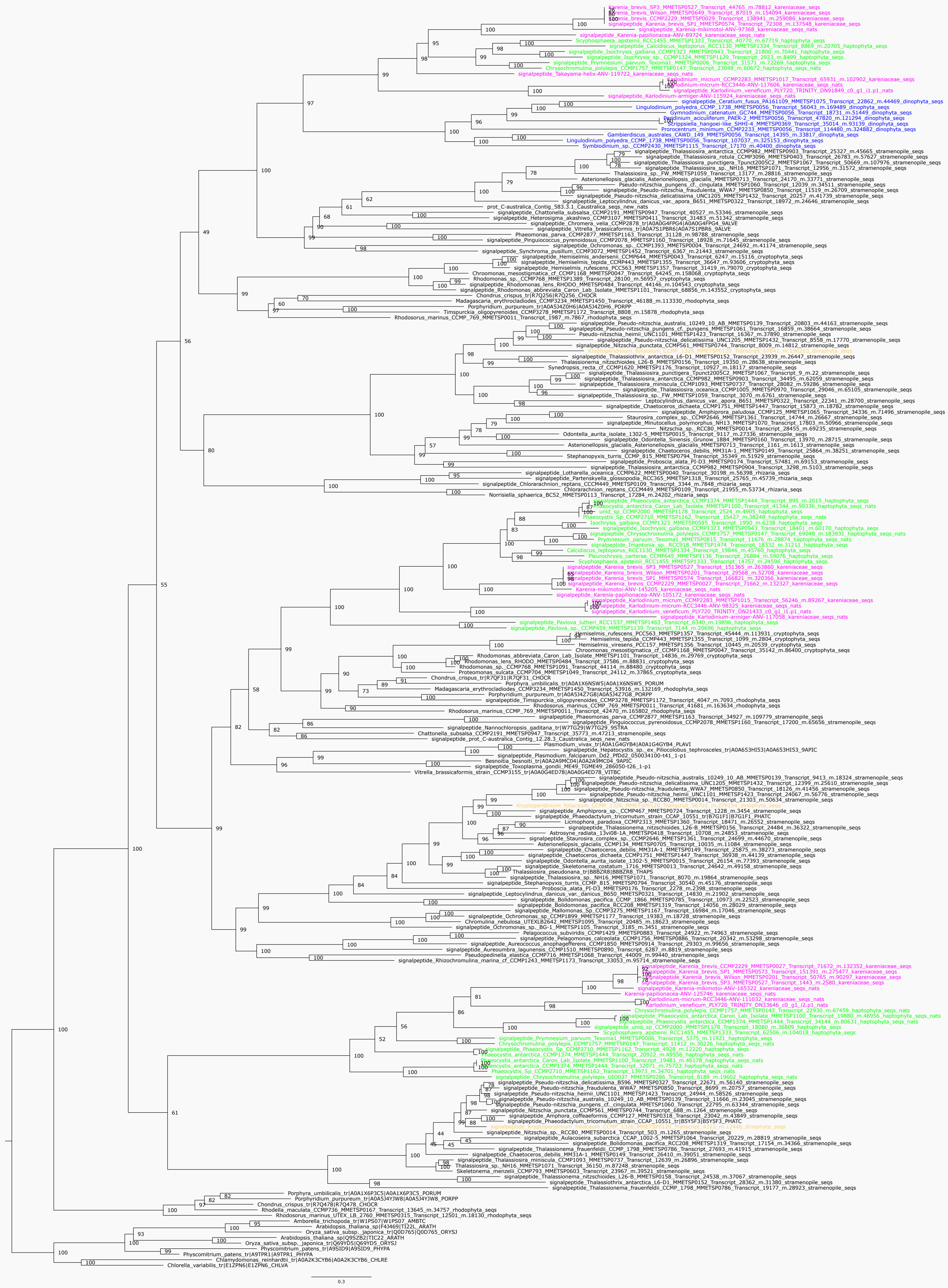

#### Tic110

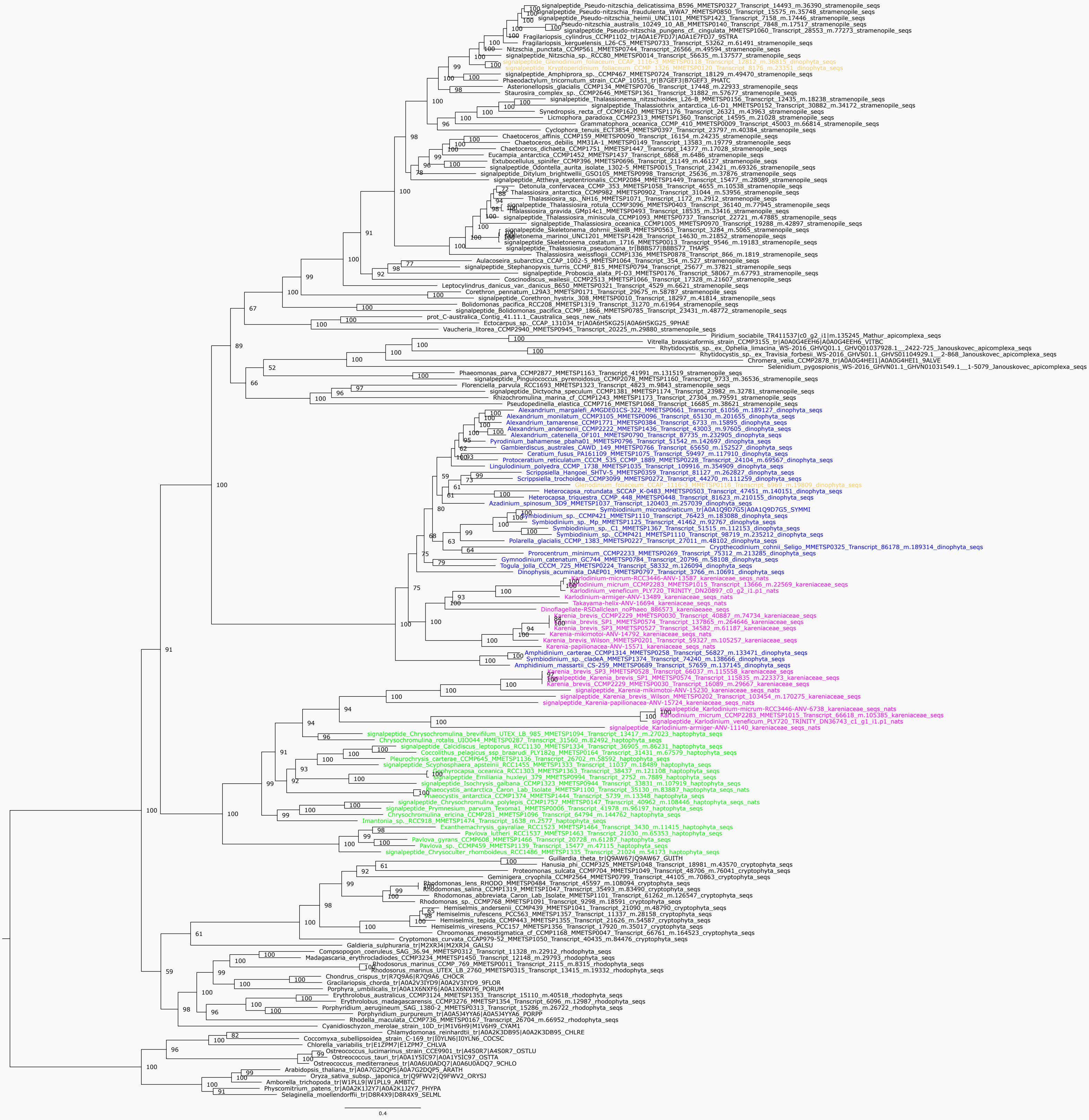

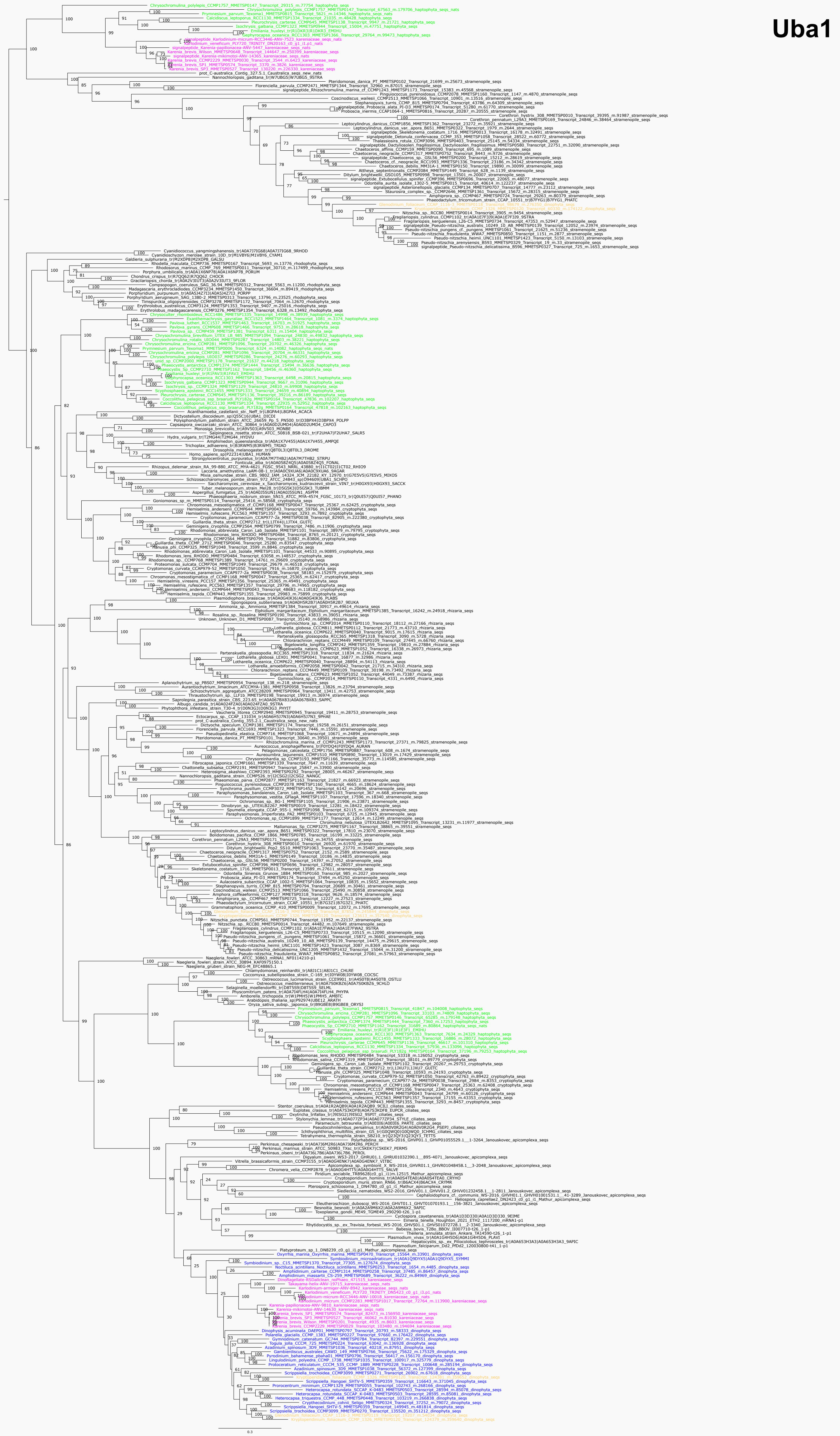

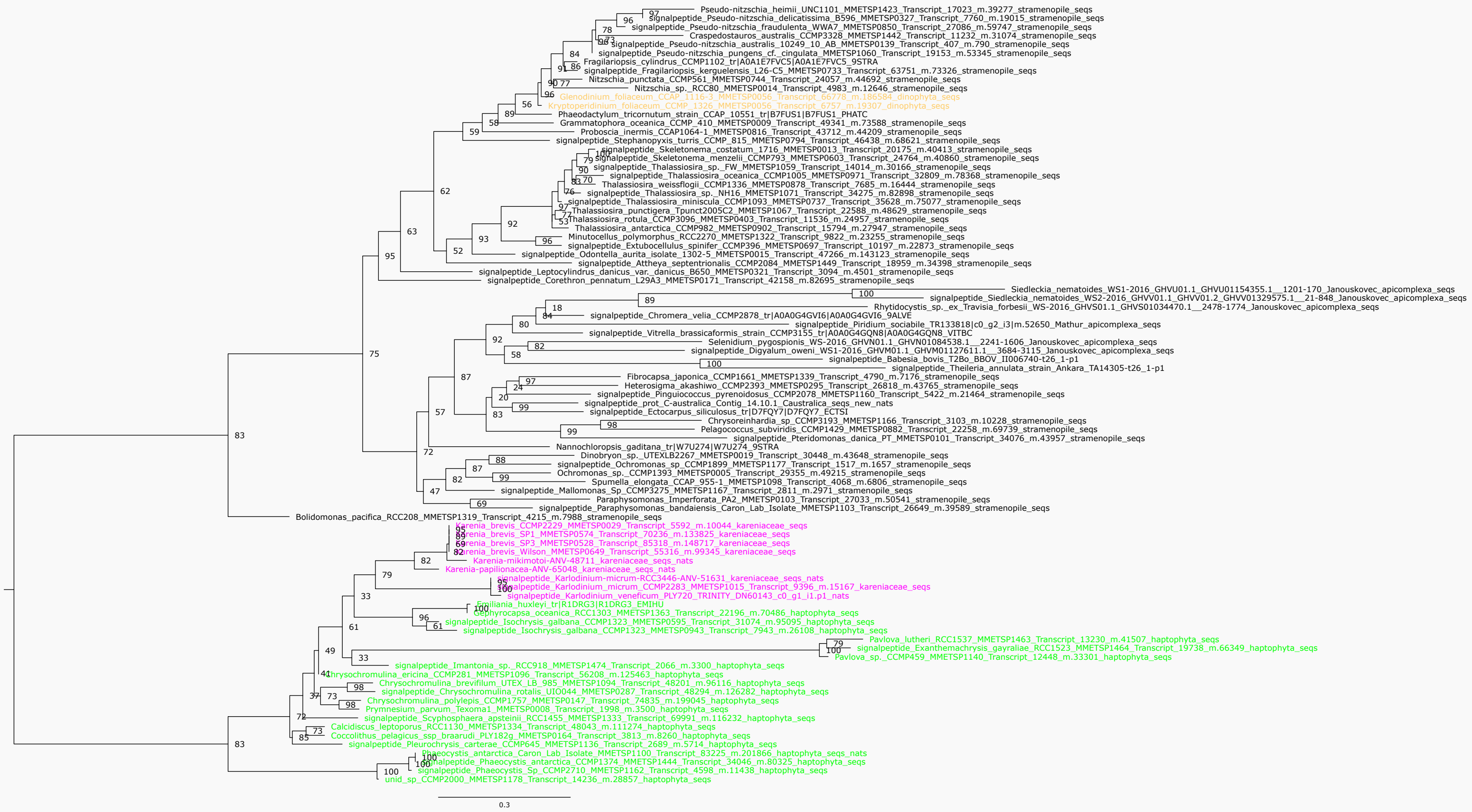

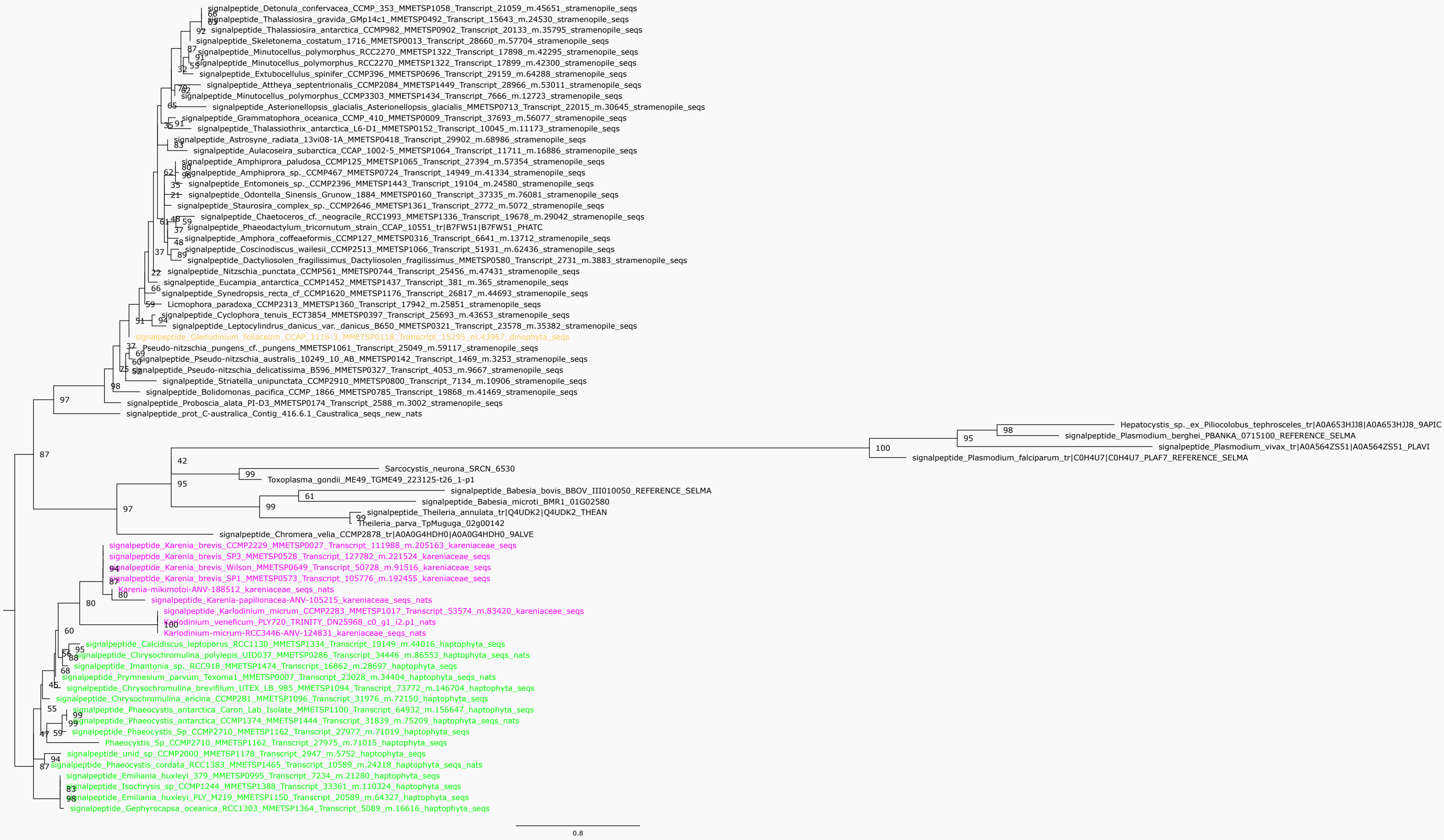
